# Standardized and reproducible measurement of decision-making in mice

**DOI:** 10.1101/2020.01.17.909838

**Authors:** The International Brain Laboratory, Valeria Aguillon-Rodriguez, Dora E. Angelaki, Hannah M. Bayer, Niccolò Bonacchi, Matteo Carandini, Fanny Cazettes, Gaelle A. Chapuis, Anne K. Churchland, Yang Dan, Eric E. J. Dewitt, Mayo Faulkner, Hamish Forrest, Laura M. Haetzel, Michael Hausser, Sonja B. Hofer, Fei Hu, Anup Khanal, Christopher S. Krasniak, Inês Laranjeira, Zachary F. Mainen, Guido T. Meijer, Nathaniel J. Miska, Thomas D. Mrsic-Flogel, Masayoshi Murakami, Jean-Paul Noel, Alejandro Pan-Vazquez, Cyrille Rossant, Joshua I. Sanders, Karolina Z. Socha, Rebecca Terry, Anne E. Urai, Hernando M. Vergara, Miles J. Wells, Christian J. Wilson, Ilana B. Witten, Lauren E. Wool, Anthony Zador

## Abstract

Progress in science requires standardized assays whose results can be readily shared, compared, and reproduced across laboratories. Reproducibility, however, has been a concern in neuroscience, particularly for measurements of mouse behavior. Here we show that a standardized task to probe decision-making in mice produces reproducible results across multiple laboratories. We designed a task for head-fixed mice that combines established assays of perceptual and value-based decision making, and we standardized training protocol and experimental hardware, software, and procedures. We trained 140 mice across seven laboratories in three countries, and we collected 5 million mouse choices into a publicly available database. Learning speed was variable across mice and laboratories, but once training was complete there were no significant differences in behavior across laboratories. Mice in different laboratories adopted similar reliance on visual stimuli, on past successes and failures, and on estimates of stimulus prior probability to guide their choices. These results reveal that a complex mouse behavior can be successfully reproduced across multiple laboratories. They establish a standard for reproducible rodent behavior, and provide an unprecedented dataset and open-access tools to study decision-making in mice. More generally, they indicate a path towards achieving reproducibility in neuroscience through collaborative open-science approaches.

## Introduction

Progress in science depends on reproducibility and thus requires standardized assays whose methods and results can be readily shared, compared, and reproduced across laboratories (Baker, 2016; Ioannidis, 2005). Such assays are common in fields such as astronomy (Fish et al., 2016; The H.E.S.S. Galactic plane survey, 2018), physics (CERN Education, Communications and Outreach Group, 2018), genetics (Dickinson et al., 2016), and medicine (Bycroft et al., 2018), and perhaps rarer in fields such as sociology (Camerer et al., 2018) and psychology (Forscher et al., 2020; Frank et al., 2017; Makel et al., 2012). They are also rare in neuroscience, a field that faces a reproducibility crisis (Baker, 2016; Botvinik-Nezer et al., 2020; Button et al., 2013).

Reproducibility has been a particular concern for measurements of mouse behavior (Kafkafi et al., 2018). Though the methods can be generally reproduced across laboratories, the results can be surprisingly different (“methods reproducibility” vs. “results reproducibility”, Goodman, Fanelli and Ioannidis, 2016). Even seemingly simple assays that probe responses to pain or stress can be swayed by extraneous factors such as the identity or sex of the experimenter. Behavioral assays can be difficult to reproduce across laboratories even when they share a similar apparatus (Chesler et al., 2002; Crabbe et al., 1999; Sorge et al., 2014; Tuttle et al., 2018).

The growing arsenal of genetic, imaging, and physiological tools available for mouse brains facilitate mechanistic studies of decision making (Carandini and Churchland, 2013; Glickfeld et al., 2014; O’Connor et al., 2009). The International Brain Laboratory is a collaboration that aims to leverage these approaches by exploring the same mouse behavior in multiple laboratories (The International Brain Laboratory, 2017). It is thus crucial that the relevant behavioral assays be reproducible both in methods and in results. Equally, it is crucial that data and techniques be made available to the broader community in an open-science approach (Beraldo et al., 2019; Charles et al., 2020; Forscher et al., 2020; Koscielny et al., 2014; Poldrack and Gorgolewski, 2014; de Vries et al., 2020).

Studying decision-making requires a task that places specific sensory, cognitive, and motor demands over hundreds of trials, affording strong constraints to behavior. The task should be complex enough to expose the neural computations that support decision-making but simple enough for mice to learn, and easily extendable to study further aspects of perception and cognition. Moreover, we sought to have ready access to the brain for neural recordings and manipulations, so we considered tasks that involve head fixation.

To meet these criteria we defined a task that combines two established assays: the classical “two-alternative forced-choice” perceptual task (Carandini and Churchland, 2013; Tanner Jr. and Swets, 1954), and a stimulus-probability manipulation analogous to the “two-armed bandit” task (Corrado et al., 2005; Fan et al., 2018; Herrnstein, 1961; Lau and Glimcher, 2005; Miller et al., 2019). A similar combination of visual detection task with probability manipulation has been used in primates to study visual selective attention (Cohen and Maunsell, 2009). In the task, mice detect the presence of a visual grating of variable contrast to their left or right, and report the perceived location with a simple movement: by turning a steering wheel (Burgess et al., 2017). The probability of stimulus appearance at the two locations is asymmetric and changes across blocks of trials. Mice thus make decisions by using both their vision and their recent experience. When the visual stimulus is evident (contrast is high), they should mostly use vision, and when the visual stimulus is ambiguous (contrast is low or zero), they should consider prior information (Whiteley and Sahani, 2008) and choose the side that has recently been more likely.

We here present results from a large cohort of mice trained in the task, demonstrating reproducible methods and reproducible results across laboratories. Mice learned the task in all laboratories, though often at a different pace. After learning they performed the task in a comparable manner, and with no significant difference across laboratories. Mice in different laboratories adopted a comparable reliance on visual stimuli, on past successes and failures, and on estimates of stimulus prior probability.

To facilitate reuse and reproducibility, we describe and share the hardware and software components and the experimental protocols. We established an open-access data architecture pipeline (International Brain Laboratory et al., 2019) and use it to release the >5 million mouse choices at data.internationalbrainlab.org. These results reveal that a complex mouse behavior can be successfully reproduced across laboratories, enabling collaborative studies of brain function in behaving mice.

## Results

To train mice consistently within and across laboratories we developed a standardized training pipeline (**Figure 1a**). First, we performed surgery to implant a headbar for head-fixation (**Appendix 1**). During the subsequent recovery period we handled the mice and weighed them daily. Following recovery, we put mice on water control and habituated them to the experimental setup (**Appendix 2**). Throughout these steps, we checked for adverse effects such as developing a cataract during surgery or pain after surgery or substantial weight loss following water control (4 mice excluded out of 210) (Guo et al., 2014).

**Figure 1.**
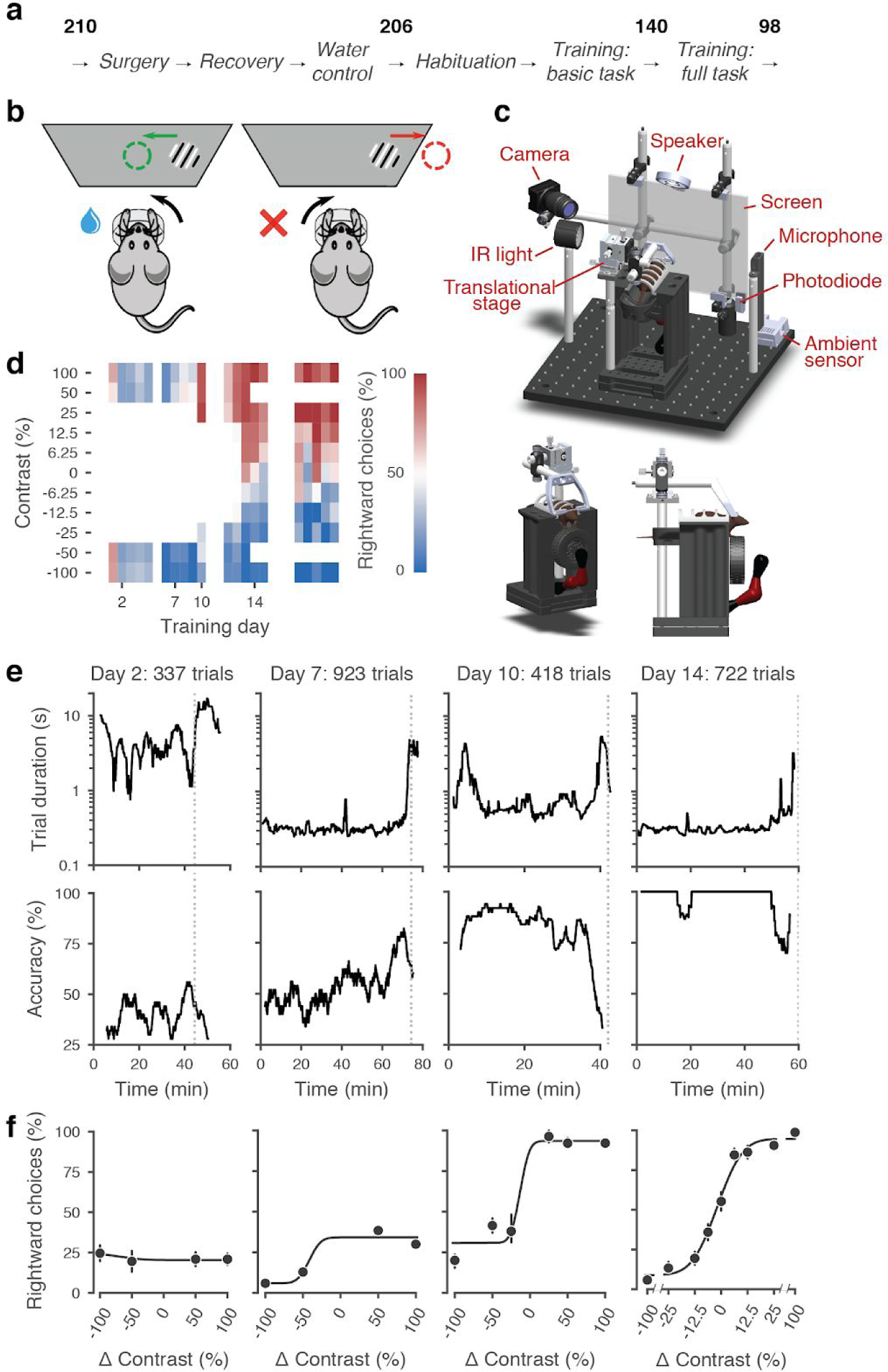
Standardized pipeline and apparatus, and training progression in the basic task. **a**, The pipeline for mouse surgeries and training. The number of animals at each stage of the pipeline is shown in bold. **b**, Schematic of the task, showing the steering wheel and the visual stimulus moving to the center of the screen vs. the opposite direction, with resulting reward vs. timeout. **c**, CAD model of the behavioral apparatus. A 3D rendered video of the CAD model can be found on [here](http://doi.org/10.6084/m9.figshare.13042574). **d**, Performance of an example mouse (KS014, from Lab 1) throughout training. Squares indicate choice performance for a given stimulus on a given day. Color indicates the percentage of right (*red*) and left (*blue*) choices. Empty squares indicate stimuli that were not presented. Negative contrasts denote stimuli on the left, positive contrasts denote stimuli on the right. **e**, Example sessions from the same mouse. *Vertical lines* indicate when the mouse reached the session-ending criteria based on trial duration (*top*) and accuracy on high-contrast trials (*bottom*) averaged over a rolling window of 10 trials. **f**, Psychometric curves for those sessions, showing the fraction of trials in which the stimulus on the right was chosen (rightward choices) as a function of stimulus position and contrast (difference between right and left, i.e. positive for right stimuli, negative for left stimuli). Circles show the mean and error bars show ± 68% confidence intervals. The training history of this mouse can be explored at [this interactive web page](http://data.internationalbrainlab.org/mouse/18a54f60-534b-4ed5-8bda-b434079b8ab8).

Mice were then trained in two stages: first they learned a *basic task*, where the probability of a stimulus appearing on the left or the right was equal (50:50), and then they learned the *full task*, where the probability of stimuli appearing on the left vs. right switched in blocks of trials between 20:80 and 80:20. Out of 206 mice that started training, 140 achieved proficiency in the basic task, and 98 in the full task (26 are still in training at the time of writing, see Figure 1 - Supplement 1d for training progression and proficiency criteria). The basic task is purely perceptual: the only informative cue is the visual stimulus. The full task instead invites integration of perception with recent experience: when stimuli are ambiguous it is best to choose the more likely option.

To facilitate reproducibility, we standardized multiple variables, measured multiple other variables, and shared the behavioral data in a database that we inspected regularly. By providing standards and guidelines, we sought to control variables such as mouse strain and provider, age range, weight range, water access, food protein and fat. We did not attempt to standardize other variables such as light-dark cycle, temperature, humidity, and environmental sound, but we documented and measured them regularly (**Figure 1 - Supplement 1a**) (Voelkl et al., 2020). Data were shared and processed across the collaboration according to a standardized web-based pipeline (International Brain Laboratory et al., 2019). This pipeline included a colony management database that stored data about each session and mouse (e.g., session start time, animal weight, etc.), a centralized data repository for files generated in the task (e.g. behavioral responses and compressed video and audio files), and a platform which provided automated analyses and daily visualizations (data.internationalbrainlab.org) (Yatsenko et al., 2018).

Mice were trained in a standardized setup involving a steering wheel placed in front of a screen (**Figure 1b,c**). The visual stimulus, a grating, appeared at variable contrast on the left or right half of the screen. The stimulus position was coupled with movements of the response wheel, and mice indicated their choices by turning the wheel left or right to bring the grating to the center of the screen (Burgess et al., 2017). Trials began after the mouse held the wheel still for 0.2-0.5 s and were announced by an auditory “go cue”. Correct decisions were rewarded with sweetened water (10% sucrose solution), whereas incorrect decisions were indicated by a noise burst and were followed by a longer inter-trial interval (2 s) (Guo et al., 2014, **Figure 1 - Supplement 2**). The experimental setups included systems for head-fixation, visual and auditory stimuli presentation, and recording of video and audio (**Figure 1c**). These were standardized and based on open-source hardware and software (**Appendix 3**).

### Training progression in the basic task

We begin by describing training in the basic task, where stimuli on the left vs. right appeared with equal probability. This version of the task is purely visual, in that no other information can be used to increase expected reward.

The training proceeded in automated steps, following predefined criteria (**Figure 1d, Appendix 2**). Initially, mice experienced only easy trials with highly visible stimuli (100% and 50% contrast). As performance improved, the stimulus set progressively grew to include contrasts of 25%, 12%, 6%, and finally 0% (**Figure 1d, Figure 1 - Supplement 1b-c**). For a typical mouse (**Figure 1d**), according to this automated schedule, stimuli with 25% contrast were introduced in training day 10, 12% contrast in Day 12, and the remaining contrasts in Day 13. On this day the 50% contrast trials were dropped, to increase the proportion of low-contrast trials. To reduce response biases, incorrect responses on easy trials (high contrast) were more likely to be followed by a “repeat trial” with the same stimulus contrast and location.

The duration of training sessions varied according to performance (**Figure 1e**). Sessions lasted at most 90 minutes and ended according to a criterion based on number of trials, total duration, and response times (**Figure 1 - Supplement 3**). For instance, for the example mouse, session on Day 2 ended when 45 min elapsed with <400 trials; sessions on Days 7, 10, and 14 ended when trial duration increased to five times above baseline (**Figure 1e**).

To encourage vigorous motor responses and to increase the number of trials, the automated protocol gradually increased the motor demands and reduced reward volume. At the beginning of training, the wheel gain was high (8 deg/mm), making the stimuli highly responsive to small wheel movements, and rewards were large (3 μL). As training progressed, the wheel gain progressively decreased to 4 deg/mm and the reward decreased to 1.5 μL, according to a predefined schedule (**Figure 1 - Supplement 1b-c**).

Mouse performance gradually improved until it provided high-quality visual psychometric curves (**Figure 1f**). At first, performance hovered around or below 50%. It could be even below 50% because of no-response trials (which were labeled as incorrect trials in our analyses), systematic response biases, and the bias-correcting procedure that tended to repeat trials following errors on easy trials (at high contrast). Performance then typically increased over days to weeks until mice made only rare mistakes (lapses) on easy trials (e.g. **Figure 1f**, Day 14).

Animals were considered to have reached proficiency in the basic task when they had been introduced to all contrast levels, and had met predefined performance criteria. These criteria required psychometric curves to satisfy minimal conditions for absolute response bias (< 16%, units of contrast), contrast sensitivity (threshold < 19%) and lapse rate (total < 0.2 on both sides) across three consecutive training sessions (**Figure 1 - Supplement 1d**). The training procedure, performance criteria, and psychometric parameters are described in detail in **Appendix 2**.

### Training succeeded but with different rates across mice and laboratories

The training procedures succeeded in all laboratories, but the duration of training varied across mice and laboratories (**Figure 2**). There was substantial variation among mice, with the fastest learners achieving basic task proficiency in 3 days, and the slowest after 59 days (**Figure 2a-c**). The average training took 18.4 ± 13.0 days (s.d., n = 140, **Figure 2b**). The number of days needed to achieve basic task proficiency was different across laboratories (**Figure 2c**, p < 0.001, Kruskal-Wallis nonparametric test followed by a post-hoc Dunn’s multiple comparisons test). Some labs had homogeneous learning rates (e.g., Lab 2 within-lab interquartile range of 8 days), while other labs had larger variability (e.g., Lab 6 interquartile range of 22 days).

**Figure 2.**
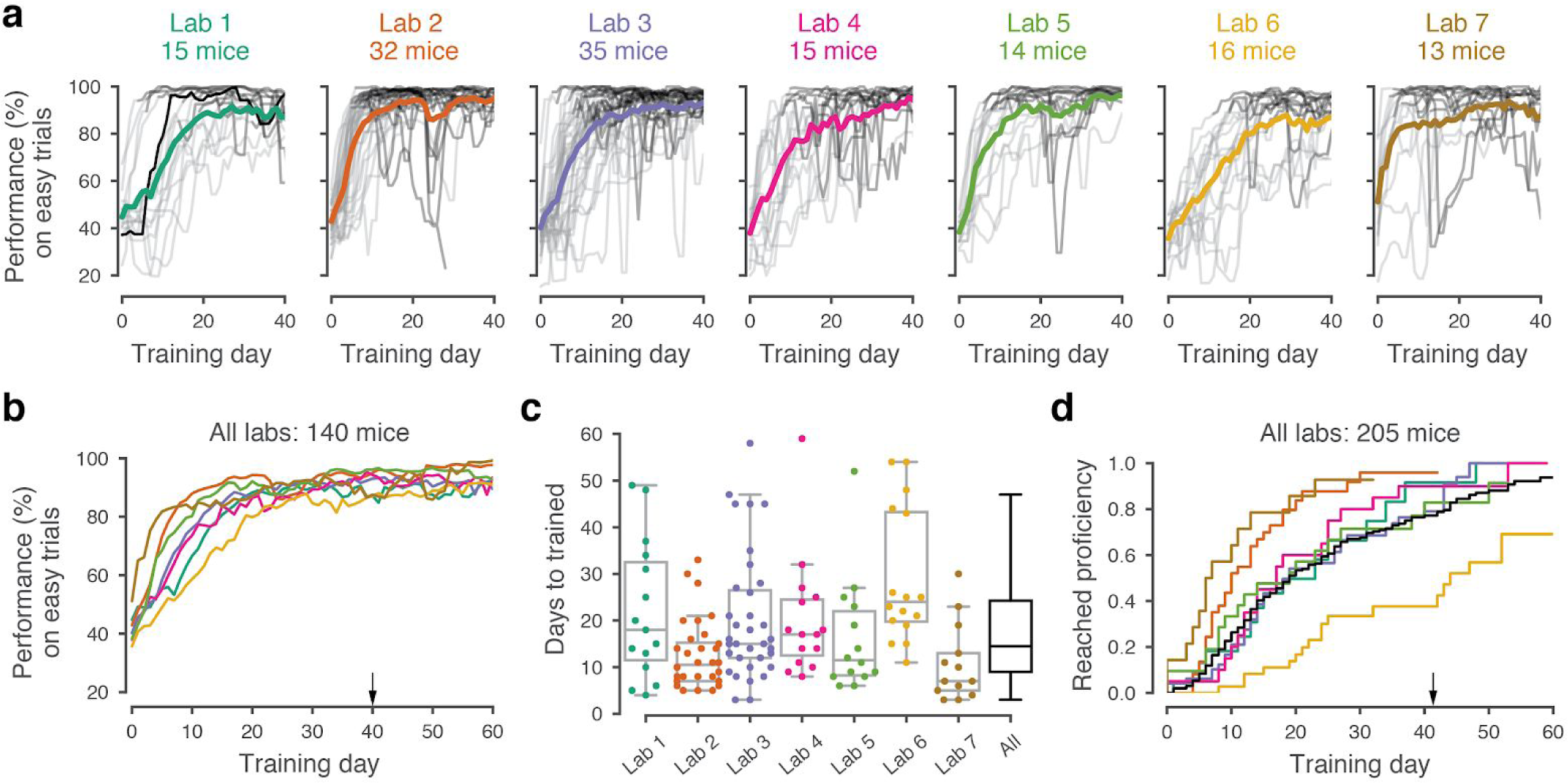
Learning rates differed across mice and laboratories. **a**, Performance curves for each mouse, for each laboratory. Performance was measured on easy trials (50% and 100% contrast). Each panel represents a different lab, and each thin curve represents a mouse. The transition from *light gray* to *dark gray* indicates when each mouse achieved proficiency in the basic task. *Black*, performance for example mouse in Fig. 1. Thick colored lines show the lab average. Curves stop at day 40, when the automated training procedure suggests that mice be dropped from the study if they have not learned. **b**, Average performance curve of each laboratory across training days. **c**, Training times for each mouse compared to the distribution across all laboratories (*black*). *Boxplots* show median and quartiles. **d**, Cumulative proportion of mice to have reached proficiency as a function of training day. *Black curve* denotes average across mice and laboratories.

The variability in performance decreased as training progressed, but it did not disappear completely. Variability in performance was larger in the middle of their training time than towards the end. For example, the variation across mice (s.d.) of performance on easy trials was 19.1% (mean 80.7%) on day 15, but 10.1% (mean 91.1%) on day 40 (**Figure 2 - Supplement 1**). Our procedures succeeded in training tens of mice in each laboratory (**Figure 2d**). Indeed, there was a 80% probability that mice would learn the task within the 40 days that were usually allotted, and when mice were trained for a longer duration, the success rate rose even further (**Figure 2d**). There was, however, variability in learning rates across mice and laboratories. This variability is intriguing and would present a challenge for projects that aim to study learning. We next ask whether the behavior of trained mice was consistent and reproducible across laboratories.

### Performance in the basic task was indistinguishable across laboratories

Once mice achieved basic task proficiency, multiple measures of performance became indistinguishable across laboratories (**Figure 3a-e**). We first examined the psychometric curves for the three sessions leading up to proficiency, which showed a stereotypical shape across mice and laboratories (**Figure 3a**). The average across mice of these psychometric curves was similar across laboratories (**Figure 3b**). The lapse rates (i.e., the errors made in response to easy contrasts of 50% and 100%) were low (9.5 ± 3.6%, **Figure 3c**) with no significant difference across laboratories. The slope of the curves, which measures contrast sensitivity, was also similar across laboratories, at 14.3 ± 3.8 (s.d., n = 7 laboratories, **Figure 3d**). Finally, the horizontal displacement of the curve, which measures response bias, was small at 0.3 ± 8.4 (s.d., n = 7, **Figure 3e**). None of these measures showed a significant difference across laboratories, either in median (performance: p = 0.63, threshold: p = 0.81, bias: p = 0.81, FDR corrected Kruskal-Wallis test) or in variance (performance: p = 0.09, threshold: p = 0.57, bias: p = 0.57, FDR corrected Levene’s test). Indeed, the within-lab consistency of choice fractions was not larger than expected by chance (**Figure 3 - Supplement 1a**).

**Figure 3.**
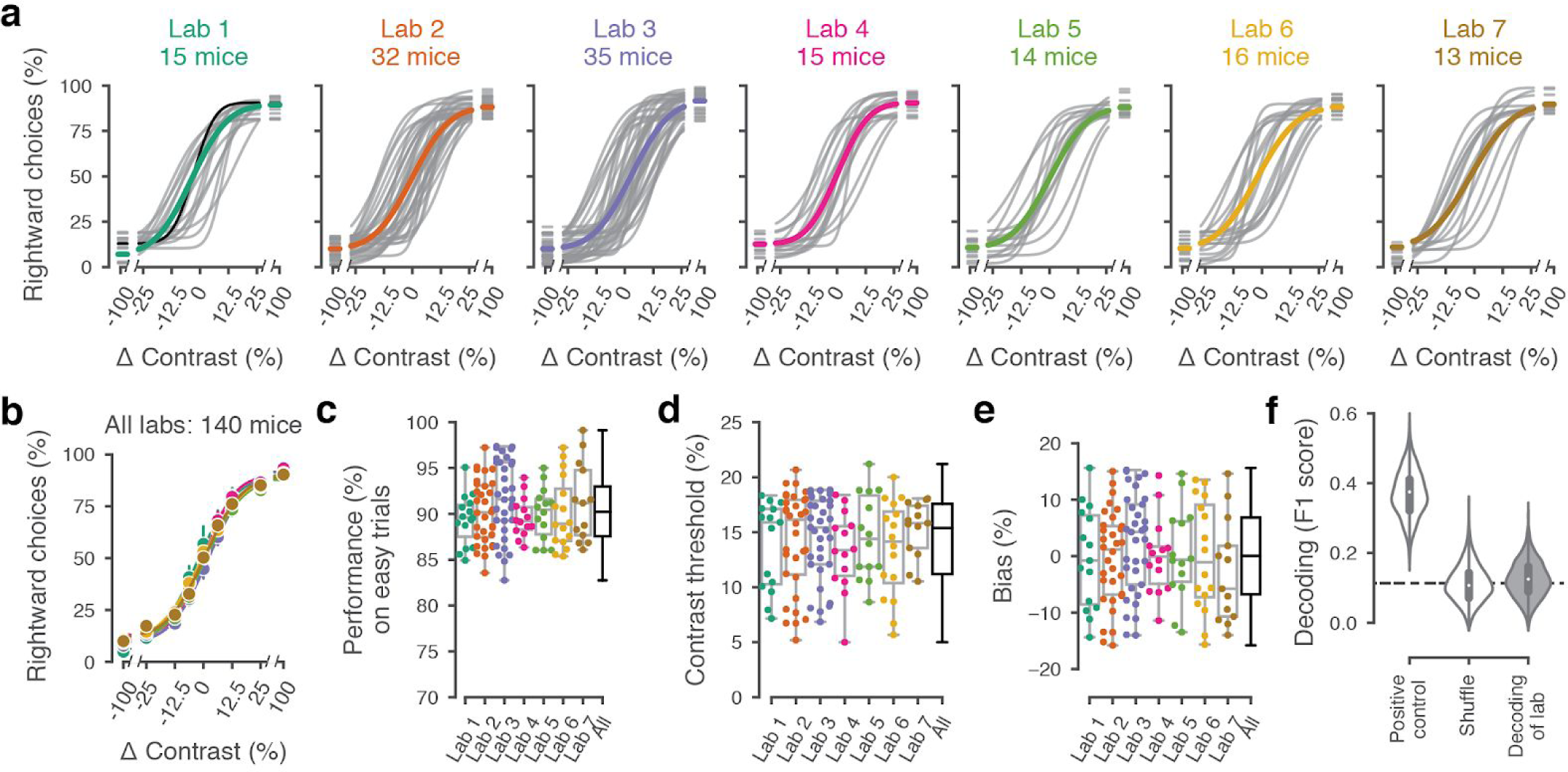
Performance in the basic task was indistinguishable across laboratories. **a**, Psychometric curves across mice and laboratories for the three sessions at which mice achieved proficiency on the basic task. Each curve represents a mouse (*gray*). Black curve represents the example mouse in Fig. 1. Thick colored lines show the lab average. **b**, Average psychometric curve for each laboratory. Circles show the mean and error bars ± 68% CI. **c**, performance on easy trials (50 and 100% contrasts) for each mouse, plotted per lab and over all labs. Colored dots show individual mice and boxplots show the median and quartiles of the distribution. **d**-**e**, Same, for visual threshold and bias. **f**, Performance of a Naive Bayes classifier trained to predict from which lab mice belonged, based on the measures in **c**-**e**. We included the timezone of the laboratory as a positive control and generated a null-distribution by shuffling the lab labels. Dashed black line represents chance-level classification performance. Violin plots show the full distribution of the 2,000 random sub-samples of 8 mice per laboratory, the white dots show the median, thick lines the interquartile range and thin lines 1.5x the interquartile range.

Variations across laboratories were also small in terms of trial duration and number of trials per session, even though no specific effort was made to harmonize these variables (**Figure 3 - Supplement 2**). The median time from stimulus onset to feedback (trial duration, a coarse measure of reaction time) was 468 ± 221 ms, showing some differences across laboratories (**Figure 3 - Supplement 2**, p = 0.004, Kruskal-Wallis nonparametric test). Mice on average performed 719 ± 223 trials per session (**Figure 3 - Supplement 2**). This difference was significant (p < 10^−6^, one-way ANOVA) but only in one laboratory relative to the rest. It may reflect different experimenter decisions on when to end training sessions: our standard protocol suggested but did not mandate when to end a session.

Similar results were observed in the behavior of the mice during subsequent sessions. We compared the behavioral performance across laboratories in the first three sessions of the full task (described later). Again we found that behavior was not significantly different across laboratories (**Figure 3 - Supplement 3**).

Variation in performance across laboratories was so low that we were not able to assign a mouse to a laboratory based on performance. Having found little variation in behavioral variables when considered one by one (**Figure 3c-e**), we next asked whether mice from different laboratories may exhibit characteristic combinations of these variables. We thus trained a Naive Bayes classifier (Pedregosa et al., 2011), with 2000 random sub-samples of 8 mice per laboratory, to predict lab membership from these behavioral variables. First, we tested whether the classifier showed the expected behavior when provided with an informative variable: the time zone in which animals were trained. In this positive control, the classifier performed above chance (**Figure 3f**, *Positive control*) and the confusion matrix showed clusters of labs which are located in the same time zone (**Figure 3 - Supplement 4c**). Next, we generated a null-distribution by shuffling the lab labels (**Figure 3f**, *Shuffle*). Finally we used the classifier to predict in which lab a mouse was trained. The classifier failed to identify the laboratory of origin of the mice: it performed at chance level (**Figure 3f**, *Decoding of lab*), with an F1 score of 0.12 ± 0.051, and a 95th percentile of the distribution included the chance level of 0.11 (mean of the null-distribution). The most common classifications were off-diagonal in the confusion matrix, and hence incorrect (**Figure 3 - Supplement 4**). Similar results were obtained with two other classifier algorithms (**Figure 3 - Supplement 4**).

We repeated this analysis for the first three sessions of the full task. Again, there was no significant difference across laboratories on the performance, visual threshold, and bias, and lab membership could not be identified above chance by classifiers (**Figure 3 - Supplement 3**). Moreover, as we will see in the next section, performance remained indistinguishable across laboratories also in the full version of the task.

### Performance in the full task was indistinguishable across laboratories

After training the mice in the purely perceptual, basic task, we introduced them to the *full task*, where optimal performance requires integration of perception with recent experience (**Figure 4a,b**). Specifically, we introduced block-wise biases in the probability of stimulus location, and therefore the more likely correct choice. Sessions started with a block of unbiased trials (50:50 probability of left vs. right) and then alternated between blocks of variable length (20-100 trials) biased towards the right (20:80 probability) or towards the left (80:20 probability) (**Figure 4a**). The transition between blocks was not signaled, so the mice had to estimate a prior for stimulus location based on recent task statistics. This task invites the mice to integrate information across trials and to use this prior knowledge in their perceptual decisions.

**Figure 4.**
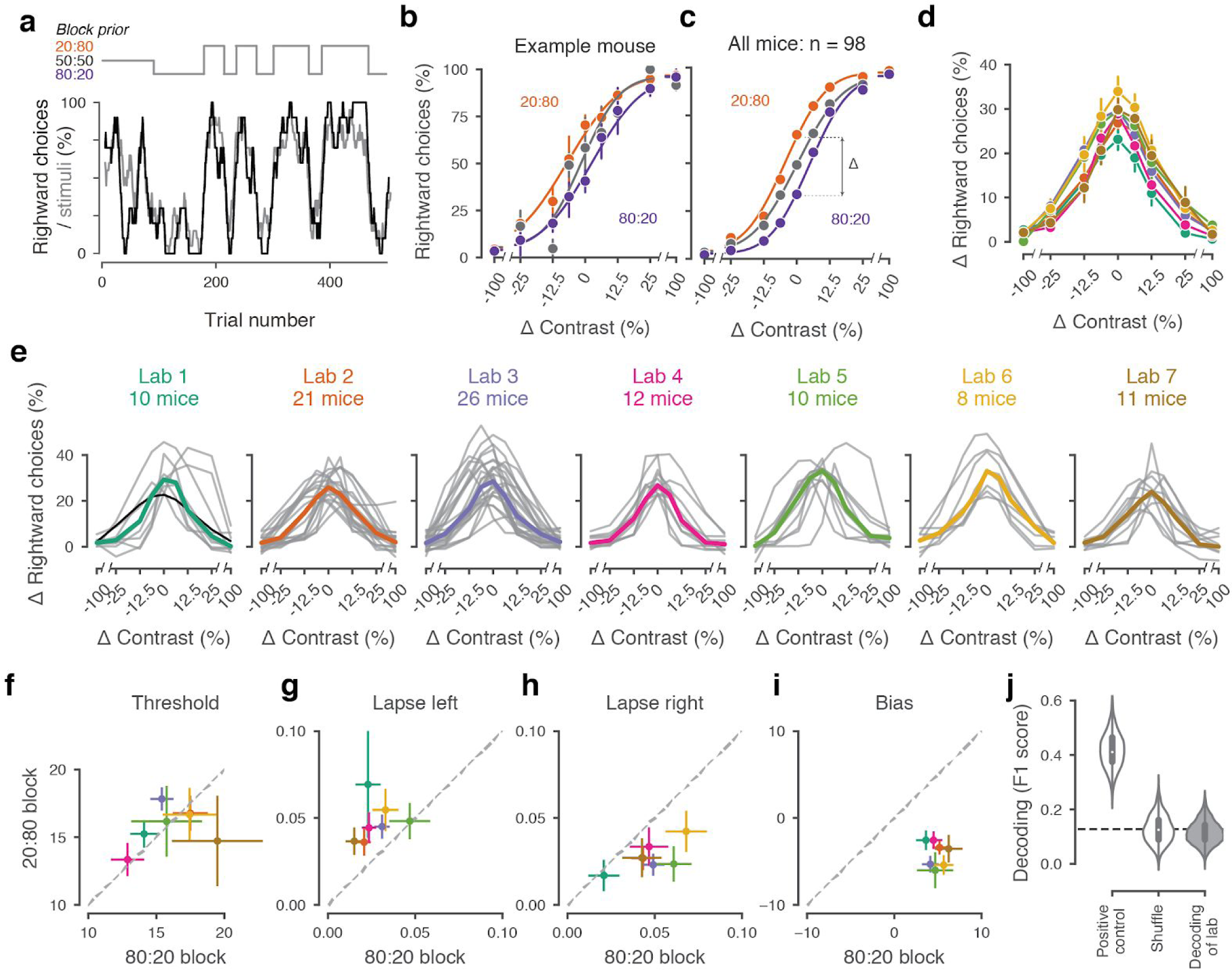
Mice successfully integrate priors into their decisions and task strategy. **a**, Block structure in an example session. Each session started with 90 trials of 50:50 prior probability, followed by alternating 20:80 and 80:20 blocks of varying length. Presented stimuli (grey, 10-trial running average) and the mouse’s choices (black, 10-trial running average) track the block structure. **b**, Psychometric curves shift between blocks for the example mouse. **c**, For each mouse that achieved proficiency on the full task (Figure 1 - Supplement 1d) and for each stimulus, we computed a ‘bias shift’ by reading out the difference in choice fraction between the 20:80 and 80:20 blocks (dashed lines). **d**, Average shift in rightward choices between block types, as a function of contrast for each laboratory (colors as in 2c, 3c; error bars show mean ± 68% CI). **e**, Shift in rightward choices as a function of contrast, separately for each lab. Each line represents an individual mouse (grey), with the example mouse in black. Thick colored lines show the lab average. **f**, Visual threshold, **g**, left lapses, **h**, right lapses, and **i**, bias separately for the 20:80 and 80:20 block types. Each lab is shown as mean +- s.e.m. **j**, Classifier results as in 3f, based on all data points in f-i.

To assess how mice used information about block structure, we compared their psychometric curves in the different block types (**Figure 4b,c**). Mice incorporated block priors into their choices, already from the first sessions in which they were exposed to this full task (**Figure 3 - Supplement 3**). To assess proficiency in the full task we used a fixed set of criteria (**Figure 1 - Supplement 1d**). We considered performance in the three sessions in which mice reached proficiency on the full task (**Figure 1 - Supplement 1d**). The curves for the 20:80 and 80:20 blocks were shifted relative to the curve for the 50:50 block, with mice more likely to choose right in the 20:80 blocks (where right stimuli appeared 80% of the time) and to choose left in the 80:20 blocks (where left stimuli appeared 80% of the time). As expected, block structure had the greatest impact on choices when sensory evidence was absent (contrast = 0%, **Figure 4c**). In these conditions, it makes sense for the mice to be guided by recent experience, and thus to choose differently depending on the block prior.

Changes in block type had a similar effect on mice in all laboratories (**Figure 4d,e**). The average shift in rightward choices was invariably highest at 0% contrast, where rightward choices in the two blocks differed by an average of 28.5%. This value did not significantly differ across laboratories any more than it differed within laboratories (one-way ANOVA F(6) = 1.345, p = 0.2455, **Figure 4d,e**).

An analysis of the psychometric curves showed highly consistent effects of block type, with no significant differences across laboratories (**Figure 4f-i**). Changes in block type did not significantly affect the visual threshold, estimated from the slope of the psychometric curve (Wilcoxon Signed-Rank test, p = 0.85, n = 97 mice, **Figure 4f**). However, it did change the lapse rates, which were consistently higher on the left in the 20:80 blocks and on the right in the 80:20 blocks (Wilcoxon Signed-Rank test, lapse left; p < 10^−6^, lapse right; p < 10^−7^, **Figure 4g,h**). Finally, as expected there was a highly consistent change in overall bias, with curves shifting to the left in 20:80 trials and to the right in 80:20 trials (Wilcoxon Signed-Rank test, p < 10^−16^, **Figure 4i**). Just as in the basic task (Figure 3), a classifier trained on these variables could not predict above chance the origin laboratory of individual mice above chance (**Figure 4j**). Moreover, neither choice fractions nor bias shifts showed within-lab consistency that was larger than expected by chance (**Figure 3 - Supplement 1b,c**). This confirms that mice performed the full task similarly across laboratories.

### A probabilistic model reveals common strategies across mice and laboratories

Lastly, we investigated the strategies used by the mice in the basic task and in the full task and asked where these strategies were similar across mice and laboratories. Mice might incorporate non-sensory information into their decisions, even when such information is not predictive of reward (Busse et al., 2011; Lak et al., 2020a). Therefore, even when reaching comparable performance levels, different mice might be weighing task variables differently when making a decision.

To quantify how different mice form their decisions, we used a simple generalized linear model (**Figure 5a**). The model is based on similar approaches used in both value-based (Lau and Glimcher, 2005) and sensory-based decision making (Busse et al., 2011; Pinto et al., 2018). In the model, the probability of making a right choice is calculated from a logistic function of the linear weighted sum of several predictors: the stimulus contrast, the outcome of the previous trial, and a bias term that represents the overall preference of a mouse for a particular choice across sessions (Busse et al., 2011). In the case of the full task we added a parameter representing the identity of the block, measuring the weight of the prior stimulus statistics in the mouse decisions (**Figure 5a**).

**Figure 5.**
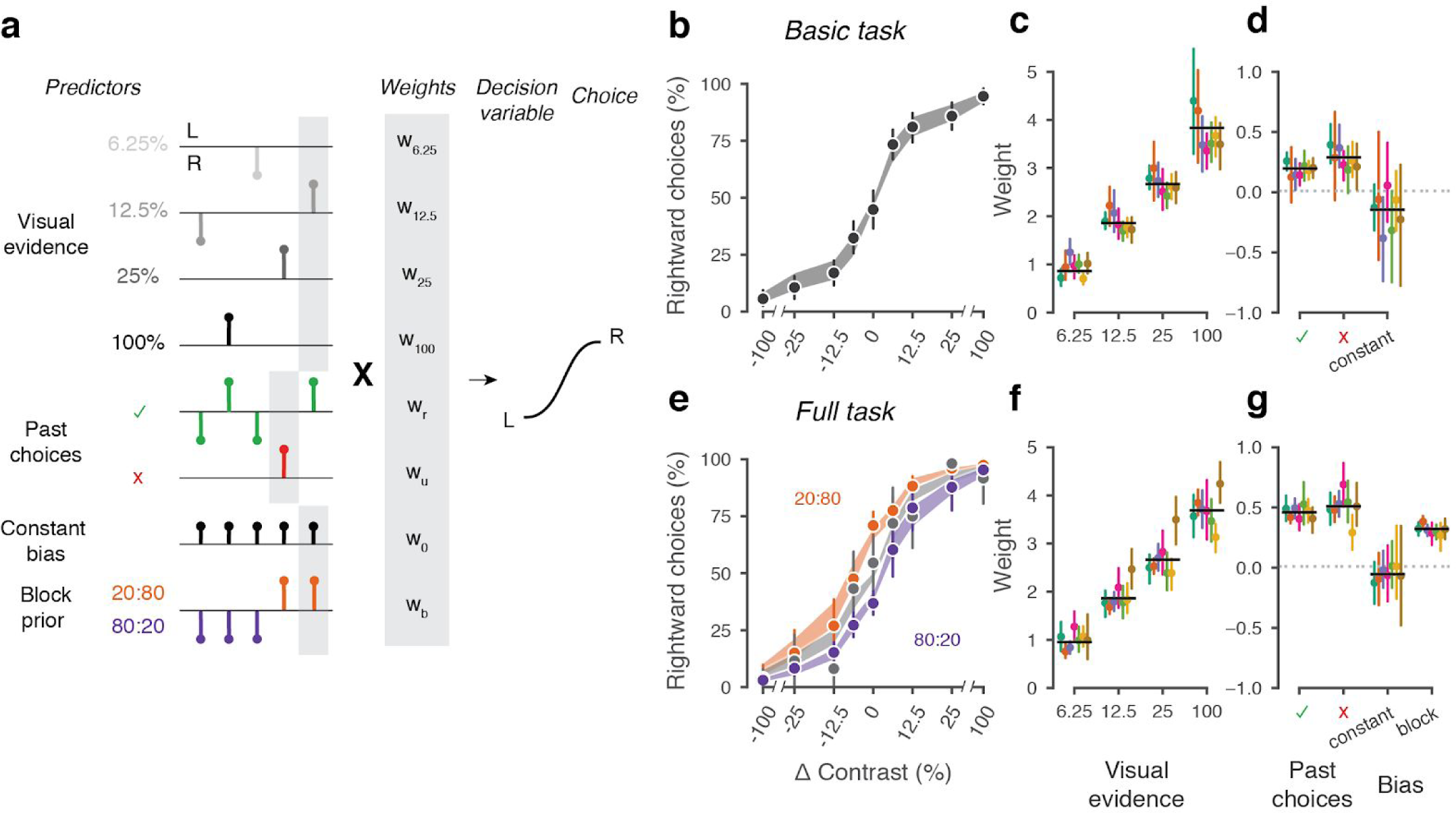
A probabilistic model reveals a common strategy across mice and laboratories. **a**, Schematic diagram of predictors included in the GLM. Each stimulus contrast (except for 0%) was included as a separate predictor. Past choices were included separately for rewarded and unrewarded trials. The dashed line represents the block prior predictor, only present in the full task. **b**, Psychometric curves from the example mouse across 3 sessions in the basic task. Shadow represents 95% confidence interval of the predicted choice fraction of the model. Points and error bars represent the mean and across-session confidence interval of the data. **c-d**, Weights for GLM predictors across labs in the basic task, error bars represent the 95% confidence interval across mice. **e-g**, as b-d but for the full task.

We fitted the model to the choices of each individual mouse over three sessions by logistic regression. The mean condition number for the basic model was 2.4 and 3.2 for the full model. The low conditions numbers for both models do not suffer from multicollinearity and the coefficients are therefore interpretable.

The model fit the mouse choices well and captured the relative importance of sensory and non-sensory information across mice and laboratories (**Figure 5b-g**). The model was able to accurately predict the behavior of individual mice, both in the basic task **(Figure 5b**) and in the full task (**Figure 5e**). As expected, visual terms had large weights, which grew with contrast to reach values above 3 at 100% contrast. Weights for non-sensory factors were much lower (**Figure 5c,f**, note different scale). Weights for past choices were positive for both rewarded (basic 0.19, full 0.42) and unrewarded previous trials (basic 0.33, full 0.46), suggesting that mice were perseverant in their choice behavior (**Figure 5d,g; Figure 5 - Supplement 2a**). Indeed, previous choices more strongly influenced behavior in the full task, both after rewarded (t(125) = 10.736, p < 10^−6^) and unrewarded trials (t(125) = 4.817, p < 10^−6^). Importantly, fitted weights and the model’s predictive accuracy (unbiased: 81.03 + 5.1%, biased 82.1 + 5.8%) (**Figure 5 - Supplement 1**) were similar across laboratories, suggesting an overall common strategy.

The model coefficients demonstrated that mice were perseverant in their actions (**Figure 5d,g; Figure 5 - Supplement 2a**). This behavior can arise from insensitivity to the outcome of previous trials and from slow drifts in the decision process across trials, arising from correlations in past choices independently of reward (Lak et al., 2020a; Mendonça et al., 2018). To disentangle these two factors, we corrected for slow across-trial drifts in the decision process (Lak et al., 2020a). This correction revealed a win-stay/lose-switch strategy in both the basic and full task, that coexists with slow drifts in choice bias across trials (**Figure 5 - Supplement 2b**). Moreover, history-dependence was modulated by confidence in the previous trial. These effects were generally consistent across laboratories.

## Discussion

These results reveal that a complex mouse behavior can be successfully reproduced across laboratories, and more generally suggest a path towards improving reproducibility in neuroscience. To study mouse behavior across laboratories we developed and implemented identical experimental equipment and a standard set of protocols. Not only did mice learn the task in all laboratories, but critically, after learning they performed the task comparably across laboratories. Mice in different laboratories had similar psychophysical performance in a purely sensory version of the task and adopted similar choice strategies in the full task, where they benefited from tracking the stimulus prior probability. Behavior showed variations across sessions and across mice, but these variations were no larger across laboratories than within laboratories.

Success did not seem guaranteed at the outset, because neuroscience faces a crisis of reproducibility (Baker, 2016; Botvinik-Nezer et al., 2020; Button et al., 2013) particularly when it comes to measurements of mouse behavior (Chesler et al., 2002; Crabbe et al., 1999; Kafkafi et al., 2018; Sorge et al., 2014; Tuttle et al., 2018). To solve this crisis, three solutions have been proposed: large studies, many teams, and upfront registration (Ioannidis, 2005). Our approach incorporates all three of these solutions. First, we collected vast amounts of data: 5 million choices from 138 mice. Second, we involved many teams, obtaining data in 7 laboratories in 3 countries. Third, we standardized the experimental protocols and data analyses upfront, which is a key component of pre-registration.

An element that may have contributed to success is the collaborative, open-science nature of our initiative (Wool and The International Brain Laboratory., 2020). Open-science collaborative approaches are increasingly taking hold in neuroscience (Beraldo et al., 2019; Charles et al., 2020; Forscher et al., 2020; Koscielny et al., 2014; Poldrack and Gorgolewski, 2014; de Vries et al., 2020). Our work benefited from collaborative development of the behavioral assay, and from frequent and regular meetings where data were reviewed across laboratories. These meetings were structured to identify problems at the origin, provide immediate feedback, and find solutions collectively. Moreover, our work benefited from constant efforts at standardization. We took great care in standardizing and documenting the behavioral apparatus and the training protocol (see Appendices), to facilitate implementation across our laboratories and to encourage wider adoption by other laboratories. The protocols, hardware designs and software code are open-source and modular, allowing adjustments to accommodate a variety of scientific questions. The data are accessible at data.internationalbrainlab.org, and include all >5 million choices made by the mice.

Another element that might have contributed to success is our choice of behavioral task, which places substantial requirements on the mice while not being too complex. Previous failures to reproduce mouse behavior across laboratories typically arose in studies of unconstrained behavior such as responses to pain or stress (Chesler et al., 2002; Crabbe et al., 1999; Kafkafi et al., 2018; Sorge et al., 2014; Tuttle et al., 2018). To be able to study decision making, and in hopes of achieving reproducibility, we designed a task that engages multiple brain processes from sensory perception and integration of evidence to combination of priors and evidence. It seems likely that reproducibility is easier to achieve if the task requirements are substantial (so there is less opportunity to engage in other behaviors) but not so complex that they fail to motivate and engage. Tasks that are too simple and unconstrained or too arbitrary and difficult may be difficult to reproduce.

There are of course multiple ways to improve on our results, e.g. by clarifying, and if desired resolving, the differences in learning rate across mice, both within and across laboratories. The learning rate is a factor that we had not attempted to control, and we cannot here ascertain the causes of its variability. We suspect that it might arise partly from variations in the expertise and familiarity of different labs with visual neuroscience and mouse behavior, which may impede standardization. If so, perhaps as experimenters gain further experience the differences in learning times will decrease. Indeed, an approach to standardizing learning rates might be to introduce full automation in behavioral training by reducing or even removing the need for human intervention (Aoki et al., 2017; Poddar et al., 2013; Scott et al., 2015). Variability in learning rates may be further reduced by individualized, dynamic training methods (Bak et al., 2016). If desired, such methods could also be aimed at obtaining uniform numbers of trials and reaction times across laboratories.

Another fruitful avenue of research would be to characterize the behavior beyond the turning of the wheel, analyzing the dynamics of the movement of the mouse limbs. The only aspect of behavior that we analyzed here is whether the wheel was turned to the left or to the right. But to turn the wheel, the mice move substantial parts of their body, and they often do so in diverse ways (e.g. some mice turn the wheel with two hands, others with one, and so on). By analyzing these movements with videography (Mathis et al., 2018) one could classify these dynamics and perhaps identify behavioral strategies that provide more insight in the performance of the task and into the diversity of behavior that we observed across sessions and across mice.

Our work could also be extended through better modeling of mouse behavior, beyond our simple model based on stimuli, choice history, and bias. The model is only a starting point for exploring the full complexity of decision computations. Indeed, we showed that ‘perseverance’ represented by the reward history weights can be confounded by slow fluctuations in decision process over trials (Lak et al., 2020a; Mendonça et al., 2018) and such history-dependence is modulated by confidence in the previous trial (Lak et al., 2020b, 2020a; Urai et al., 2017). In addition to these analyses, our data are also ideally suited to investigate other phenomena a, such as the origin of lapses (Ashwood et al., 2019; Pisupati et al., 2019), how mice track the changes between prior blocks (Norton et al., 2019), the dependence of psychophysical behavior on fluctuating engagement states (McGinley et al., 2015), and the dynamics of trial-by-trial learning within a session (Roy et al., 2020). While beyond the scope of this investigation, we hope that the availability of this large, curated dataset will serve as a benchmark for testing these and other models of decision-making.

To encourage and support community adoption of this task, we provide detailed methods and protocols. These methods and protocols provide a path to training mice in this task and reproducing their behavior across laboratories, but of course we cannot claim that they constitute the optimal path. In designing our methods, we made many choices that were based on intuition rather than rigorous experimentation. We don’t know what is crucial in these methods and what is not.

The reproducibility of this mouse behavior makes it a good candidate for studies of the brain mechanisms underlying decision making. We hope that these resources catalyze the development of new adaptations and variations of our approach, and accelerate the use of mice in high quality, reproducible studies of neural correlates of decision-making.

## Acknowledgments

We thank Charu Reddy for helping develop animal welfare and surgical procedures; George Bekheet, Filipe Carvalho, Paulo Carriço, Robb Barrett and Del Halpin for help with hardware design; Luigi Acerbi and Zoe Ashwood for advice about model fitting; and Peter Dayan and Karel Svoboda for comments on the manuscript. AEU is supported by the German National Academy of Sciences Leopoldina. LEW is supported by a Marie Skłodowska-Curie Actions fellowship. FC was supported by an EMBO long term fellowship and an AXA postdoctoral fellowship. HMV was supported by an EMBO long term fellowship. MC holds the GlaxoSmithKline / Fight for Sight Chair in Visual Neuroscience. This work was supported by grants from the Wellcome Trust (209558 and 216324) and the Simons Foundation.

## Competing Interests

J.I.S. is the owner of Sanworks LLC which provides hardware and consulting for the experimental set-up described in this work.

## Methods

All procedures and experiments were carried out in accordance with the local laws and following approval by the relevant institutions: the Animal Welfare Ethical Review Body of University College London; the Institutional Animal Care and Use Committees of Cold Spring Harbor Laboratory, Princeton University, and University of California at Berkeley; the University Animal Welfare Committee of New York University; and the Portuguese Veterinary General Board.

### Animals

Animals (all female and male C57BL6/J mice aged 3-7 months obtained from Jackson Laboratory or Charles River) were co-housed whenever possible, with a minimum enrichment of nesting material and a mouse house. Mice were kept in a 12-h light-dark cycle, and fed with food that was 5-6% fat and 18-20% protein. See **Suppl. Table 1** for details on standardization.

### Surgery

A detailed account of the surgical methods is in **Appendix 1**. Briefly, mice were anesthetized with isoflurane and head-fixed in a stereotaxic frame. The hair was then removed from their scalp, much of the scalp and underlying periosteum was removed and bregma and lambda were marked. Then the head was positioned such that there was a 0 degree angle between bregma and lambda in all directions. The headbar was then placed in one of three stereotactically defined locations and cemented in place. The exposed skull was then covered with cement and clear UV curing glue, ensuring that the remaining scalp was unable to retract from the implant.

### Materials and Apparatus

For detailed parts lists and installation instructions, see **Appendix 3**. Briefly, all labs installed standardized behavioral rigs consisting of an LCD screen (LP097QX1, LG), a custom 3D-printed mouse holder and head bar fixation clamp to hold a mouse such that its forepaws rest on a steering wheel (86652 & 32019, LEGO) (Burgess et al., 2017). Silicone tubing controlled by a pinch valve (225P011-21, NResearch) was used to deliver water rewards to the mouse. The general structure of the rig was constructed from Thorlabs parts and was placed inside an acoustical cabinet (9U acoustic wall cabinet 600 × 600, Orion). LCD screen refresh times were captured with a Bpod Frame2TTL (Sanworks). Ambient temperature, humidity and barometric air pressure were measured with the Bpod Ambient module (Sanworks), wheel position was monitored with a rotary encoder (05.2400.1122.1024, Kubler) connected to a Bpod Rotary Encoder Module (Sanworks). Video of the mouse was recorded with a USB camera (CM3-U3-13Y3M-CS, Point Grey). A speaker (HPD-40N16PET00-32, Peerless by Tymphany) was used to play task-related sounds, and an ultrasonic microphone (Ultramic UM200K, Dodotronic) was used to record ambient noise from the rig. All task-related data was coordinated by a Bpod State Machine (Sanworks). The task logic was programmed in Python and the visual stimulus presentation and video capture was handled by Bonsai (Lopes et al., 2015) and the Bonsai package BonVision (Lopes et al., 2020).

### Habituation, Training and Experimental Protocol

For a detailed protocol on animal training, see **Appendix 2**. Mice were water restricted or given access to citric acid water on weekends (Urai et al., 2020). Mice were handled for at least 10 minutes and given water in hand for at least two consecutive days prior to head fixation. On the second of these days, mice were also allowed to freely explore the rig for 10 minutes. Subsequently, mice were gradually habituated to head fixation over three consecutive days (15-20, 20-40, and 60 minutes, respectively), observing an association between the visual grating and the reward location. On each trial, with the steering wheel locked, mice passively viewed a Gabor stimulus (100% contrast, 0.1 cycles/degree spatial frequency, random phase, vertical orientation) presented on a small screen (size: approx. 246 mm diagonal active display area). The screen was positioned 8 cm in front of the animal and centralized relative to the position of eyes to cover ∼102 visual degree azimuth. The stimulus appeared for ∼10 s randomly presented at -35° (left), +35° (right), or 0° (center) and the mouse received a reward in the latter case (3µl water with 10% sucrose).

On the fourth day, the steering wheel was unlocked and coupled to the movement of the stimulus. For each trial, the mouse must use the wheel to move the stimulus from its initial location to the center to receive a reward. Initially, the stimulus moved 8°/mm of movement at the wheel surface. If the mouse completed at least 200 correct response trials within a session, the gain of the wheel for all future sessions was halved, remaining at 4°/mm. At the beginning of each trial, the mouse was required to not move the wheel for a quiescence period of 200-500 ms (randomly drawn from an exponential distribution with a mean of 350 ms). If the wheel moved during this period, the timer was reset. After the quiescence period, the stimulus appeared on either the left (−35°) or right (+35°) with a contrast randomly selected from a predefined set (initially, 50% and 100%). Simultaneously, an onset tone (5 kHz sine wave, 10 ms ramp) was played for 100 ms. As soon as the stimulus appeared, the mouse had 60 s to move the stimulus. If it correctly moved the stimulus 35° to the center of the screen, it received a 3 μL reward; if it incorrectly moved the stimulus 35° away from the center (20° visible and the rest off-screen), it received an error timeout. If the mouse responded incorrectly or failed to reach either threshold within the 60-s window, a noise burst was played for 500ms and the inter-trial interval was set to 2 s. If the response was incorrect and the contrast was ‘easy’ (≥50%), a ‘repeat’ trial followed, in which the previous stimulus contrast and location was presented with a high probability (see **Appendix 2**).

Once an animal was proficient in the basic task, it proceeded to the full task. Here, the trial structure was identical, except that stimuli were more likely to reappear on the same side for variable blocks of trials, and counterbiasing ‘repeat’ trials were not used. Each session began with 90 trials in which stimuli were equally likely to appear on the left or right (10 repetitions at each contrast), after which the probability of the stimulus appearing on the left alternated between 0.8 and 0.2 for a given block. The number of trials in each block was drawn from a truncated exponential (range = 20-100, mean 50, τ = 60)

### Classification of laboratory membership

Three different classifiers were used to try to predict in which laboratory a mouse was trained based on behavioral metrics: Naive Bayes, Random Forest, and Logistic Regression. We used the scikit-learn implementation available in Python with default configuration settings for the three classifiers. Some labs trained more mice than others resulting in an imbalanced dataset. This imbalance was corrected by taking a random subsample of 8 mice from each laboratory 2,000 times. The size of the subsample was chosen because 8 mice was the lowest number of mice over all different datasets for which classification was performed. For each random subsample, lab membership was classified using leave-one-out cross-validation. Furthermore, a null-distribution was generated by shuffling the lab labels for each subsample and classifying the shuffled data. The classification accuracy was calculated as the F1 score (equation 1) which is a standard way of measuring a classifier’s accuracy. An F1 score of 0 indicates complete misclassification and a score of 1 indicates perfect classification.

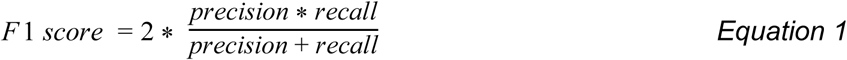

### Probabilistic choice model

To quantify and describe the different factors affecting choices across labs, we adapted a probabilistic choice model (Busse et al., 2011) used in a similar task. The model is a binomial logistic regression model, where the observer estimates the probability of choosing right (*p*) or left (1 − *p*) from sensory and non-sensory information. In the model, probabilities are obtained from the logistic transformation of the decision variable *z* (Equation 2), which in itself is the result of a weighted linear function of different task predictors (Equation 3 and Equation 4).

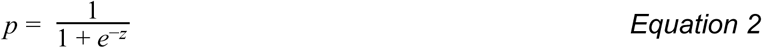

For the *basic task*, in each trial *i*, the decision variable *z* is calculated by:

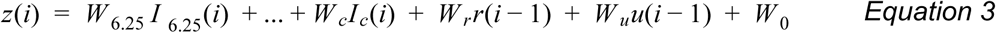

where *W*_*c*_ is is the coefficient associated with the contrast *c* ∈ {6.25, 12.5, 25, 50, 100}), and*I*_*c*_ (*i*) is an indicator function indicating +1 if the contrast *c* appeared on the right in trial *i*, -1 if it appeared on the left and 0 if that contrast was not presented. The coefficients (*W*_*r*_ and *W*_*u*_) weigh the effect of previous choices, depending on their outcome. *r*(*i* − 1) is defined as +1 when the previous trial was on the right and rewarded, -1 when on the left and rewarded, and 0 when unrewarded. Conversely, *u*(*i* − 1) is defined as +1 when the previous trial was on the right and unrewarded, -1 when on the left and unrewarded and 0 when rewarded. *W* _0_ is a constant representing the overall bias of the mouse.

When modelling the *full task* we also included the term *W*_*b*_*b*(*i*) (Equation 4), which captures the block identity *b*(*i*) for trial *i*. *b*(*i*) is defined as +1 if trial *i* is part of an 20:80 block, -1 if part of a 80:20 block and 0 if part of a 50/50 block:

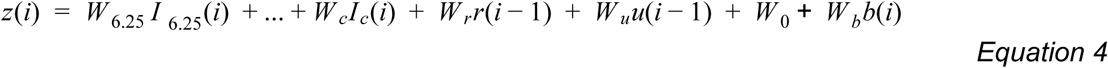

For each animal, the design matrix was built using patsy (Smith et al., 2018). The model was then fitted by regularized maximum likelihood estimation, using the *Logit*.*fit_regularized* function implemented in statsmodels (Seabold and Perktold, 2010). For the example animal (Figure 5b,e), 10.000 samples were drawn from a multivariate Gaussian (obtained from the inverse of the model’s Hessian), and for each sample the model ‘s choice fraction for each contrast level was predicted. Confidence intervals were then defined as the 0.025 and 0.975 quantiles across samples.

## Data and code availability

All data presented in this paper is publicly available. It can be viewed and accessed in two ways: via DataJoint and web browser tools at data.internationalbrainlab.org.

All data were analyzed and visualized in Python, using numpy (Harris et al., 2020), pandas (Reback et al., 2020) and seaborn (Waskom et al., 2017). Code to produce all the figures is available at github.com/int-brain-lab/paper-behavior, and a Jupyter notebook for re-creating Figure 2 can be found at jupyterhub.internationalbrainlab.org/ (under ‘resources’).

## Appendices

Appendices are available online:

Appendix 1: IBL protocol for headbar implant surgery in mice

Appendix 2: IBL protocol for mice training

Appendix 3: IBL protocol for setting up the behavioral training rig

CAD models and technical drawings for the manufactured parts can be found online. The whole CAD assembly model of the rig (including non-manufactured parts) can also be found online. A 3D rendered video of the behavioral training rig CAD model can be found online.

## Author contributions

### Task force description

The production of all IBL Platform Papers is led by a Task Force, which defines the scope and composition of the paper, assigns and/or performs the required work for the paper, and ensures that the paper is completed in a timely fashion. The Task Force members for this platform paper are Gaelle A. Chapuis, Guido T. Meijer, Alejandro Pan Vazquez, Anne E. Urai, Miles Wells, and Matteo Carandini.

### Contribution diagram

The following diagram illustrates the contributions of each author, based on the CRediT taxonomy (Brand et al., 2015). For each type of contribution there are three levels: support, equal, and lead, indicated by color in the diagram. The full written statement includes explanatory comments for each

**Supplement 1.**
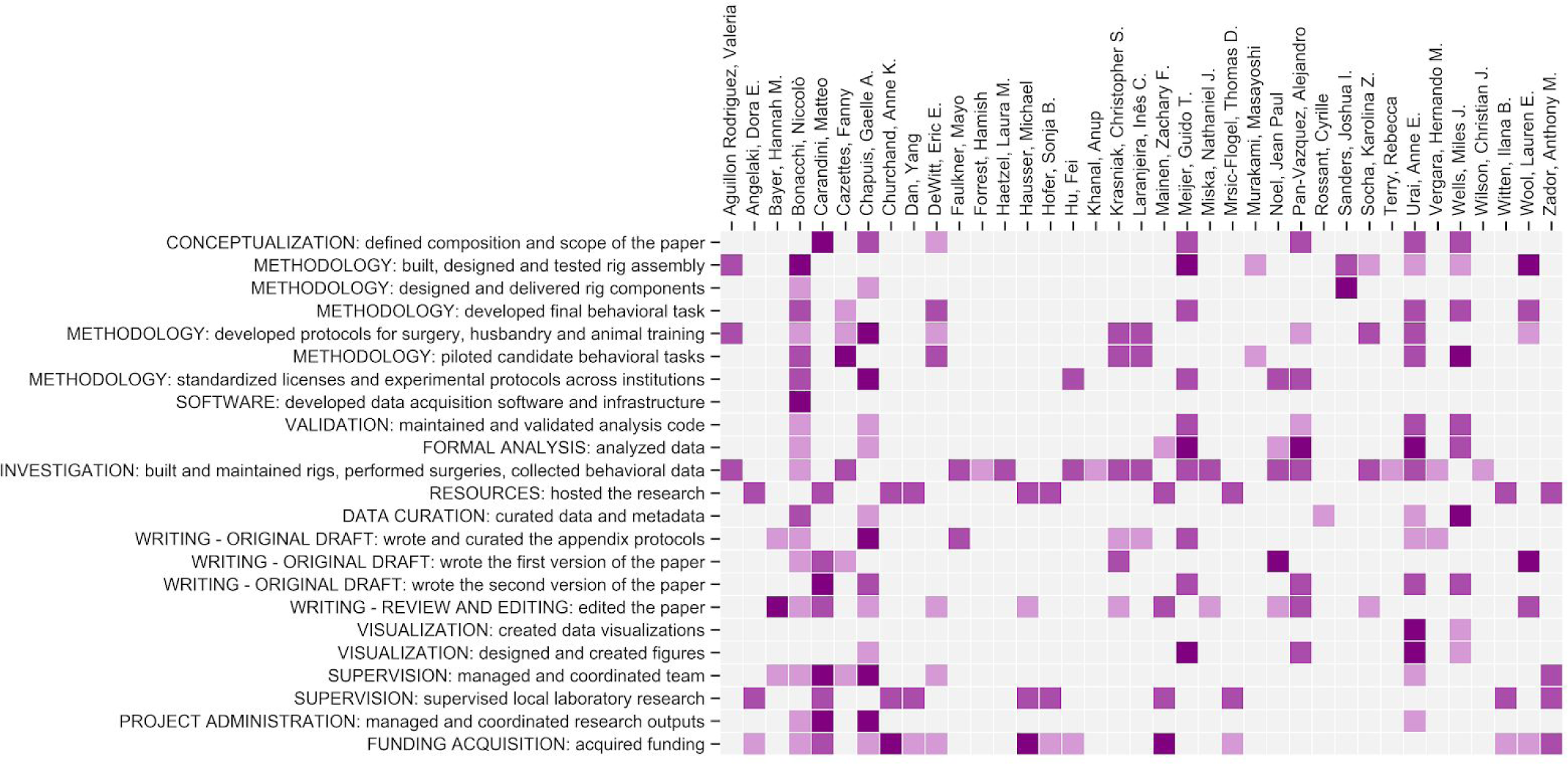
Author contributions, listed using CRediT taxonomy categories. Purple squares indicate a contribution, with levels ‘support’ (light), ‘equal’ (medium) and ‘lead’ (dark). See below for the full description of each author’s contribution.

### Contribution statement

Valeria Aguillon Rodriguez: METHODOLOGY: built, designed and tested rig assembly (equal); METHODOLOGY: developed protocols for surgery, husbandry and animal training (equal); INVESTIGATION: built and maintained rigs, performed surgeries, collected behavioral data (equal)

Dora E. Angelaki: RESOURCES: hosted the research (equal); SUPERVISION: supervised local laboratory research (equal); FUNDING ACQUISITION: acquired funding (support)

Hannah M. Bayer: WRITING - ORIGINAL DRAFT: wrote and curated the appendix protocols (support); WRITING - REVIEW AND EDITING: edited the paper (lead); SUPERVISION: managed and coordinated team (support)

Niccolò Bonacchi: METHODOLOGY: built, designed and tested rig assembly (lead); METHODOLOGY: designed and delivered rig components (support); METHODOLOGY: piloted candidate behavioral tasks (equal); METHODOLOGY: developed final behavioral task (equal); METHODOLOGY: developed protocols for surgery, husbandry and animal training (support); METHODOLOGY: standardized licenses and experimental protocols across institutions (equal); SOFTWARE: developed data acquisition software and infrastructure (lead); VALIDATION: maintained and validated analysis code (support); FORMAL ANALYSIS: analyzed data (support); INVESTIGATION: built and maintained rigs, performed surgeries, collected behavioral data (support); DATA CURATION: curated data and metadata (equal); WRITING - ORIGINAL DRAFT: wrote the first version of the paper (support); WRITING - ORIGINAL DRAFT: wrote and curated the appendix protocols (support); WRITING - REVIEW AND EDITING: edited the paper (support); SUPERVISION: managed and coordinated team (support); PROJECT ADMINISTRATION: managed and coordinated research outputs (support); FUNDING ACQUISITION: acquired funding (support)

Matteo Carandini: CONCEPTUALIZATION: defined composition and scope of the paper (lead); RESOURCES: hosted the research (equal); WRITING - ORIGINAL DRAFT: wrote the first version of the paper (equal); WRITING - ORIGINAL DRAFT: wrote the second version of the paper (lead); WRITING - REVIEW AND EDITING: edited the paper (equal); SUPERVISION: supervised local laboratory research (equal); SUPERVISION: managed and coordinated team (lead); PROJECT ADMINISTRATION: managed and coordinated research outputs (lead); FUNDING ACQUISITION: acquired funding (equal)

Fanny Cazettes: METHODOLOGY: piloted candidate behavioral tasks (lead); METHODOLOGY: developed final behavioral task (support); METHODOLOGY: developed protocols for surgery, husbandry and animal training (support); INVESTIGATION: built and maintained rigs, performed surgeries, collected behavioral data (equal); WRITING - ORIGINAL DRAFT: wrote the first version of the paper (support); SUPERVISION: managed and coordinated team (support)

Gaelle A. Chapuis: CONCEPTUALIZATION: defined composition and scope of the paper (equal); METHODOLOGY: developed protocols for surgery, husbandry and animal training (lead); METHODOLOGY: designed and delivered rig components (support); METHODOLOGY: standardized licenses and experimental protocols across institutions (lead); VALIDATION: maintained and validated analysis code (support); FORMAL ANALYSIS: analyzed data (support); DATA CURATION: curated data and metadata (support); WRITING - ORIGINAL DRAFT: wrote the second version of the paper (equal); WRITING - ORIGINAL DRAFT: wrote and curated the appendix protocols (lead); WRITING - REVIEW AND EDITING: edited the paper (support); VISUALIZATION: designed and created figures (support); SUPERVISION: managed and coordinated team (lead); PROJECT ADMINISTRATION: managed and coordinated research outputs (lead); FUNDING ACQUISITION: acquired funding (support)

Anne K. Churchand: RESOURCES: hosted the research (equal); SUPERVISION: supervised local laboratory research (equal); FUNDING ACQUISITION: acquired funding (lead)

Yang Dan: RESOURCES: hosted the research (equal); SUPERVISION: supervised local laboratory research (equal); FUNDING ACQUISITION: acquired funding (support)

Eric E. DeWitt: CONCEPTUALIZATION: defined composition and scope of the paper (support); METHODOLOGY: developed final behavioral task (equal); METHODOLOGY: piloted candidate behavioral tasks (equal); METHODOLOGY: developed protocols for surgery, husbandry and animal training (support); WRITING - REVIEW AND EDITING: edited the paper (support); SUPERVISION: managed and coordinated team (support); FUNDING ACQUISITION: acquired funding (support)

Mayo Faulkner: INVESTIGATION: built and maintained rigs, performed surgeries, collected behavioral data (equal); WRITING - ORIGINAL DRAFT: wrote and curated the appendix protocols (equal)

Hamish Forrest: INVESTIGATION: built and maintained rigs, performed surgeries, collected behavioral data (support) Laura M. Haetzel: INVESTIGATION: built and maintained rigs, performed surgeries, collected behavioral data (equal)

Michael Hausser: RESOURCES: hosted the research (equal); WRITING - REVIEW AND EDITING: edited the paper (support); SUPERVISION: supervised local laboratory research (equal); FUNDING ACQUISITION: acquired funding (lead)

Sonja B. Hofer: RESOURCES: hosted the research (equal); SUPERVISION: supervised local laboratory research (equal); FUNDING ACQUISITION: acquired funding (support)

Fei Hu: METHODOLOGY: standardized licenses and experimental protocols across institutions (equal); INVESTIGATION: built and maintained rigs, performed surgeries, collected behavioral data (equal); FUNDING ACQUISITION: acquired funding (support)

Anup Khanal: INVESTIGATION: built and maintained rigs, performed surgeries, collected behavioral data (support)

Christopher S. Krasniak: METHODOLOGY: piloted candidate behavioral tasks (equal); METHODOLOGY: developed protocols for surgery, husbandry and animal training (equal); INVESTIGATION: built and maintained rigs, performed surgeries, collected behavioral data (equal); WRITING - ORIGINAL DRAFT: wrote the first version of the paper (equal); WRITING - ORIGINAL DRAFT: wrote and curated the appendix protocols (support); WRITING - REVIEW AND EDITING: edited the paper (support)

Inês C. Laranjeira: METHODOLOGY: piloted candidate behavioral tasks (equal); METHODOLOGY: developed protocols for surgery, husbandry and animal training (equal); INVESTIGATION: built and maintained rigs, performed surgeries, collected behavioral data (equal); WRITING - ORIGINAL DRAFT: wrote and curated the appendix protocols (support)

Zachary F. Mainen: FORMAL ANALYSIS: analyzed data (support); RESOURCES: hosted the research (equal); WRITING - REVIEW AND EDITING: edited the paper (equal); SUPERVISION: supervised local laboratory research (equal); FUNDING ACQUISITION: acquired funding (lead)

Guido T. Meijer: CONCEPTUALIZATION: defined composition and scope of the paper (equal); METHODOLOGY: developed final behavioral task (equal); METHODOLOGY: built, designed and tested rig assembly (lead); METHODOLOGY: standardized licenses and experimental protocols across institutions (equal); VALIDATION: maintained and validated analysis code (equal); FORMAL ANALYSIS: analyzed data (lead); INVESTIGATION: built and maintained rigs, performed surgeries, collected behavioral data (equal); WRITING - ORIGINAL DRAFT: wrote the second version of the paper (equal); WRITING - ORIGINAL DRAFT: wrote and curated the appendix protocols (equal); VISUALIZATION: designed and created figures (lead)

Nathaniel J. Miska: INVESTIGATION: built and maintained rigs, performed surgeries, collected behavioral data (equal); WRITING - REVIEW AND EDITING: edited the paper (support)

Thomas D. Mrsic-Flogel: RESOURCES: hosted the research (equal); SUPERVISION: supervised local laboratory research (equal); FUNDING ACQUISITION: acquired funding (support)

Masayoshi Murakami: METHODOLOGY: built, designed and tested rig assembly (support); METHODOLOGY: piloted candidate behavioral tasks (support)

Jean Paul Noel: METHODOLOGY: standardized licenses and experimental protocols across institutions (equal); FORMAL ANALYSIS: analyzed data (support); INVESTIGATION: built and maintained rigs, performed surgeries, collected behavioral data (equal); WRITING - ORIGINAL DRAFT: wrote the first version of the paper (lead); WRITING - REVIEW AND EDITING: edited the paper (support)

Alejandro Pan-Vazquez: CONCEPTUALIZATION: defined composition and scope of the paper (equal); METHODOLOGY: standardized licenses and experimental protocols across institutions (equal); METHODOLOGY: developed protocols for surgery, husbandry and animal training (support); VALIDATION: maintained and validated analysis code (support); FORMAL ANALYSIS: analyzed data (lead); INVESTIGATION: built and maintained rigs, performed surgeries, collected behavioral data (equal); WRITING - ORIGINAL DRAFT: wrote the second version of the paper (equal); WRITING - REVIEW AND EDITING: edited the paper (equal); VISUALIZATION: designed and created figures (equal)

Cyrille Rossant: DATA CURATION: curated data and metadata (support)

Joshua I. Sanders: METHODOLOGY: designed and delivered rig components (lead); METHODOLOGY: built, designed and tested rig assembly (equal)

Karolina Z. Socha: METHODOLOGY: developed protocols for surgery, husbandry and animal training (equal); METHODOLOGY: built, designed and tested rig assembly (support); INVESTIGATION: built and maintained rigs, performed surgeries, collected behavioral data (equal); WRITING - REVIEW AND EDITING: edited the paper (support)

Rebecca Terry: INVESTIGATION: built and maintained rigs, performed surgeries, collected behavioral data (support)

Anne E. Urai: CONCEPTUALIZATION: defined composition and scope of the paper (equal); METHODOLOGY: built, designed and tested rig assembly (support); METHODOLOGY: piloted candidate behavioral tasks (equal); METHODOLOGY: developed final behavioral task (equal); METHODOLOGY: developed protocols for surgery, husbandry and animal training (equal); VALIDATION: maintained and validated analysis code (equal); FORMAL ANALYSIS: analyzed data (lead); INVESTIGATION: built and maintained rigs, performed surgeries, collected behavioral data (equal); DATA CURATION: curated data and metadata (support); WRITING - ORIGINAL DRAFT: wrote the second version of the paper (equal); WRITING - ORIGINAL DRAFT: wrote and curated the appendix protocols (support); VISUALIZATION: designed and created figures (lead); VISUALIZATION: created data visualizations (lead); SUPERVISION: managed and coordinated team (support); PROJECT ADMINISTRATION: managed and coordinated research outputs (support)

Hernando M. Vergara: INVESTIGATION: built and maintained rigs, performed surgeries, collected behavioral data (support); WRITING - ORIGINAL DRAFT: wrote and curated the appendix protocols (support)

Miles J. Wells: CONCEPTUALIZATION: defined composition and scope of the paper (equal); METHODOLOGY: built, designed and tested rig assembly (support); METHODOLOGY: piloted candidate behavioral tasks (lead); METHODOLOGY: developed final behavioral task (equal); VALIDATION: maintained and validated analysis code (equal); FORMAL ANALYSIS: analyzed data (equal);

DATA CURATION: curated data and metadata (lead); WRITING - ORIGINAL DRAFT: wrote the second version of the paper (equal); VISUALIZATION: designed and created figures (support); VISUALIZATION: created data visualizations (support)

Christian J. Wilson: INVESTIGATION: built and maintained rigs, performed surgeries, collected behavioral data (support)

Ilana B. Witten: RESOURCES: hosted the research (equal); SUPERVISION: supervised local laboratory research (equal); FUNDING ACQUISITION: acquired funding (support)

Lauren E. Wool: METHODOLOGY: built, designed and tested rig assembly (lead); METHODOLOGY: developed final behavioral task (equal); METHODOLOGY: developed protocols for surgery, husbandry and animal training (support); WRITING - ORIGINAL DRAFT: wrote the first version of the paper (lead); WRITING - REVIEW AND EDITING: edited the paper (equal); FUNDING ACQUISITION: acquired funding (support)

Anthony M. Zador: RESOURCES: hosted the research (equal); SUPERVISION: supervised local laboratory research (equal); SUPERVISION: managed and coordinated team (equal); FUNDING ACQUISITION: acquired funding (equal)

## Supplement

**Figure 1 - Supplement 1.**
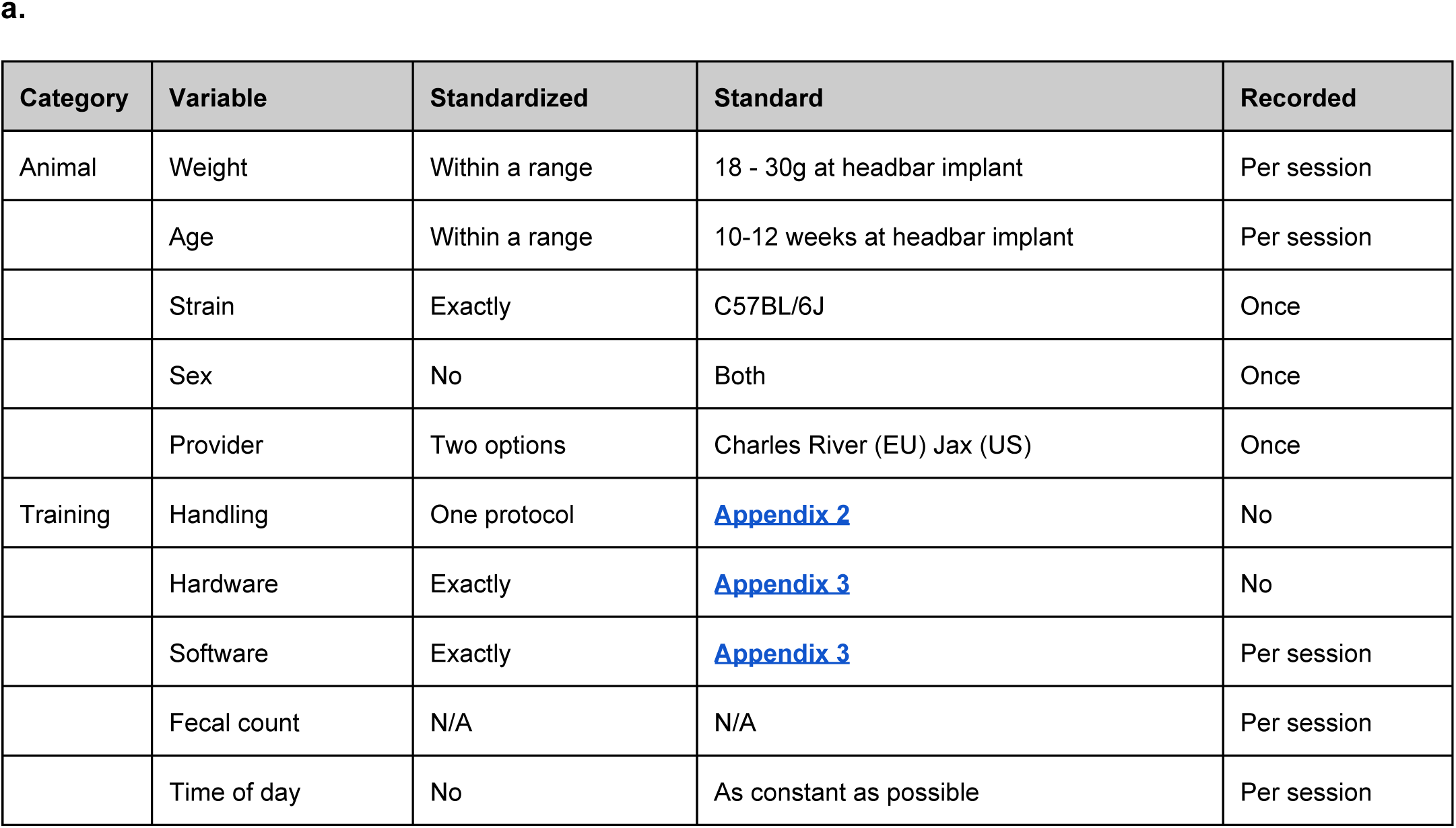
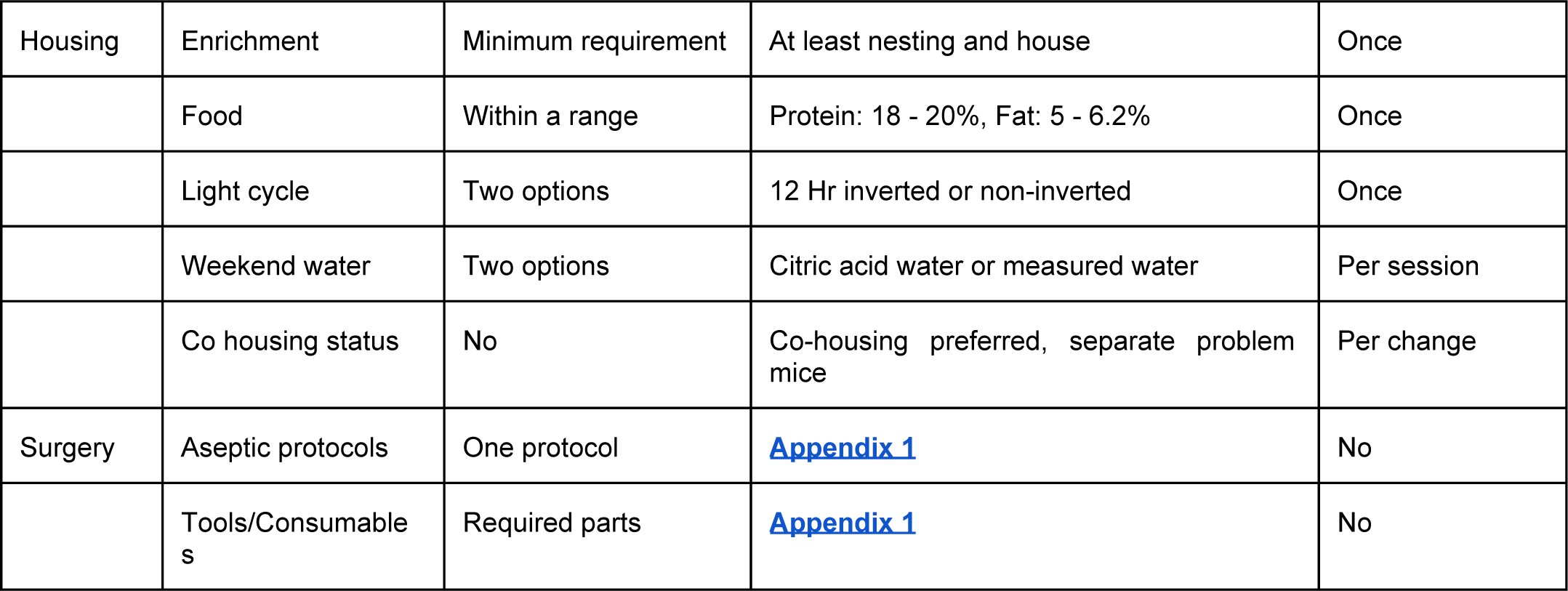
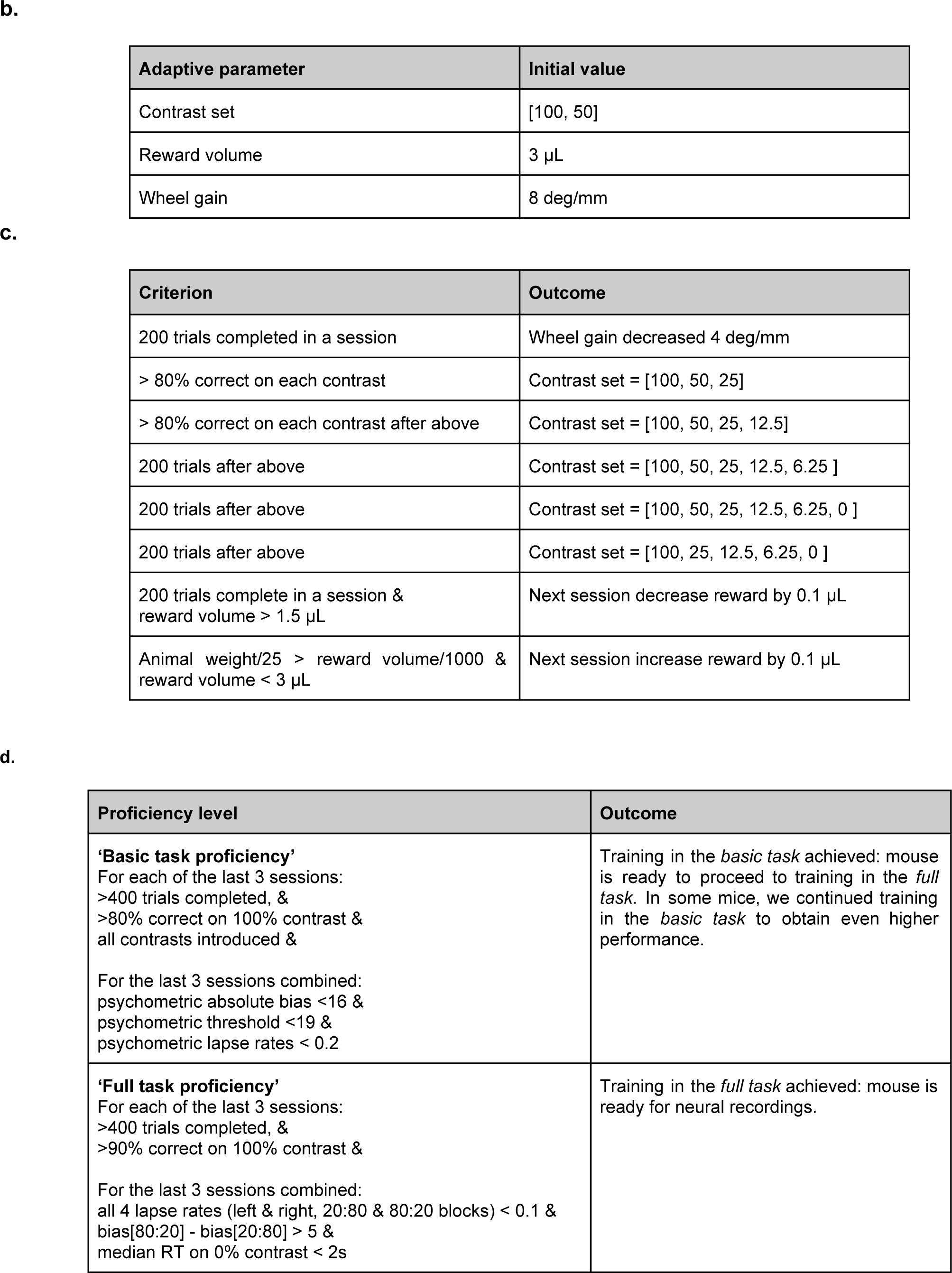
Standardization. **(a)** To facilitate reproducibility we standardized multiple aspects of the experiment. Some variables were kept strictly the same across mice, while others were kept within a range or simply recorded (see ‘Standardized’ column). **(b-c)** The behavior training protocol was also standardized. **(b)** Several task parameters adaptively changed within or across sessions **(b)** contingent on various performance criteria being met, including number of trials completed, amount of water received and proportion of correct responses.

**Figure 1 - Supplement 2.**
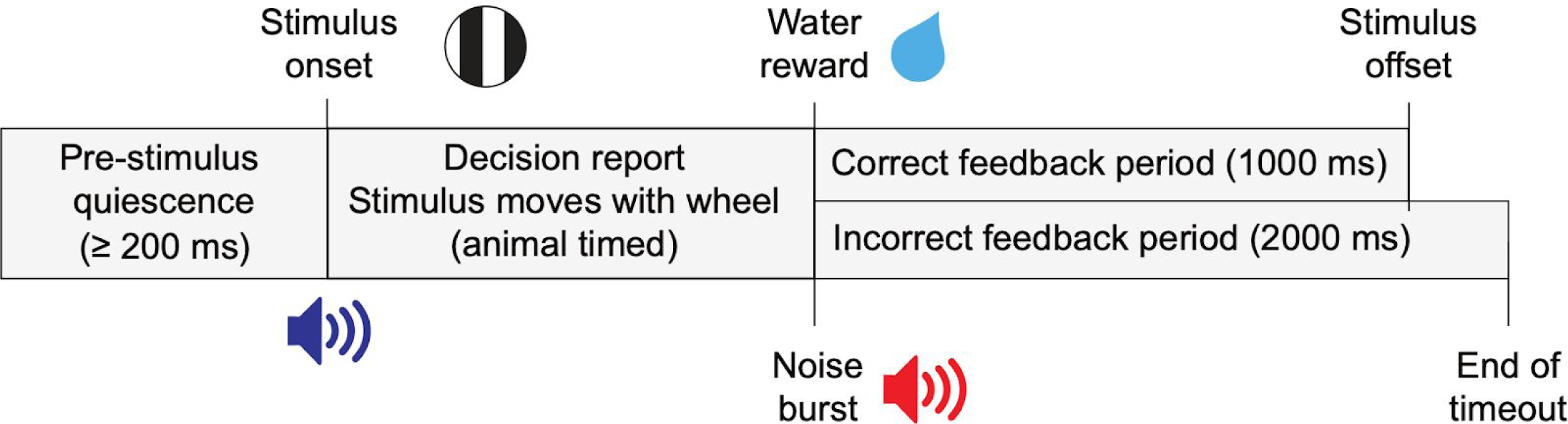
Task trial structure. Trials began with an enforced quiescent period during which the wheel must be kept still for at least 200ms, after which was the visual stimulus onset and an audio tone to indicate the start of the closed-loop period. The feedback period began when a response was given or 60s had elapsed since stimulus onset. On correct trials a reward was given and the stimulus remained in the center of the screen for 1s. On incorrect trials there was a noise burst and a 2s timeout before the next trial.

**Figure 1 - Supplement 3:**
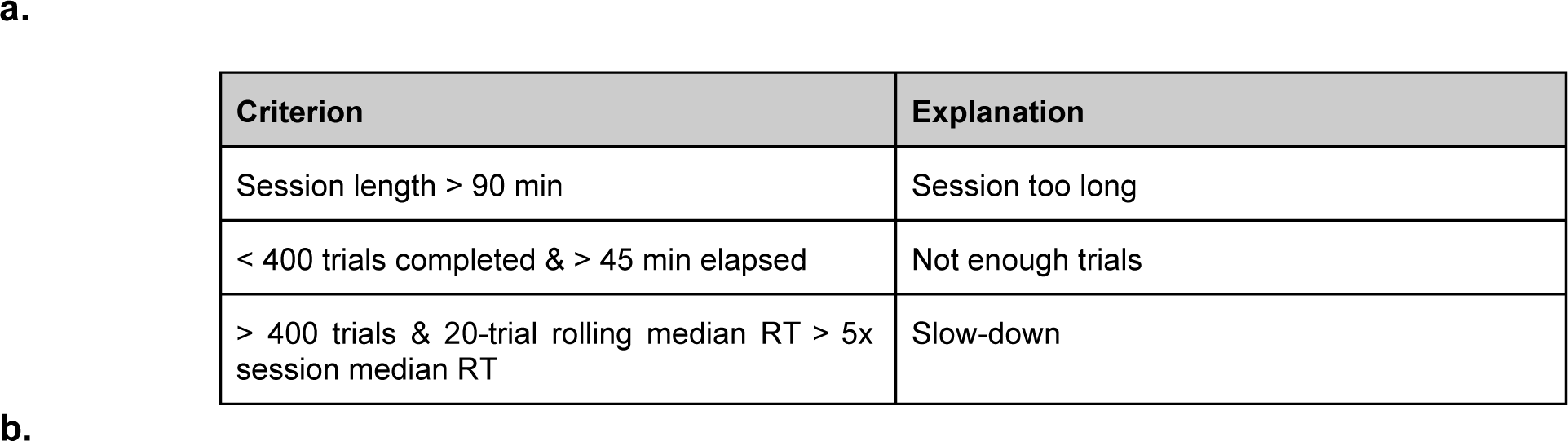
Within-session disengagement criteria.

**Figure 1 - Supplement 4.**
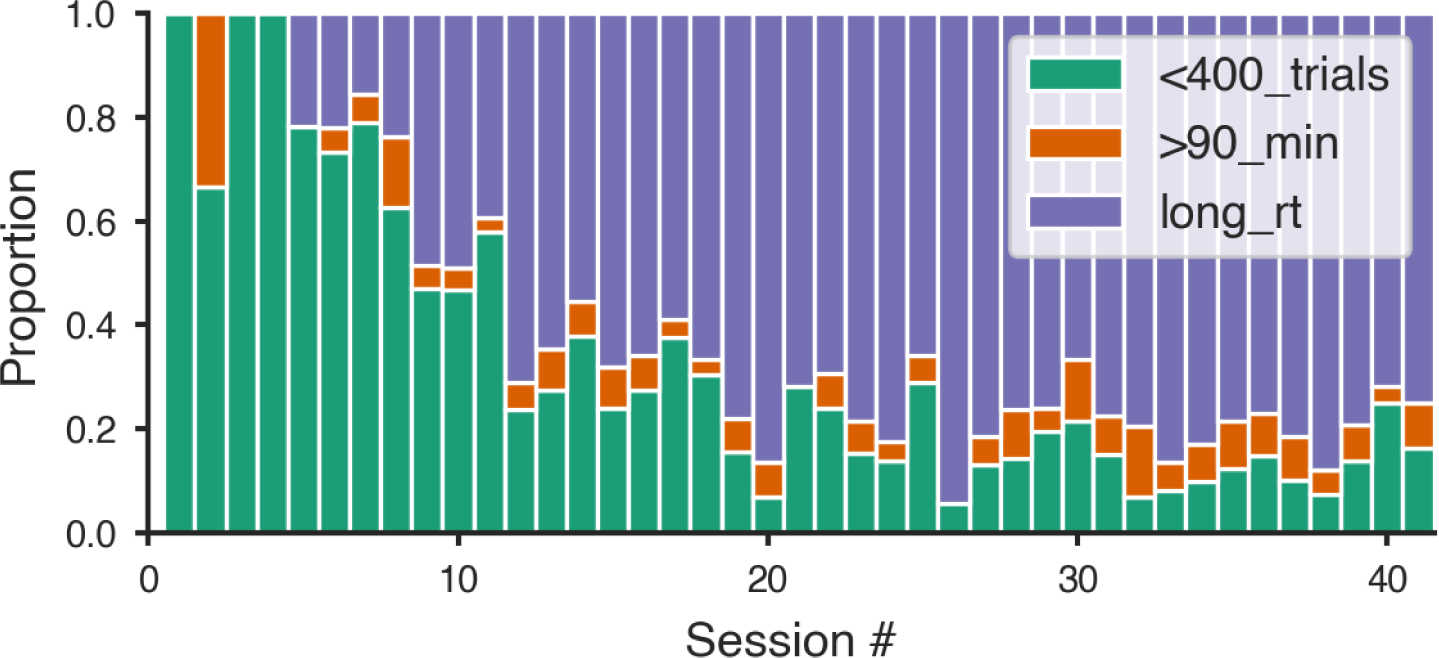
Distribution of within-session disengagement criteria. **(b)** Proportion of sessions that ended in each of the 3 criteria for all mice that learned the task. (**a**) The three criteria were, 1. Fewer than 400 trials in 45 minutes (green); 2. over 400 trials performed and median reaction time over the last 20 trials was over 5x the median for the whole session (blue); 3. Over 400 trials performed and session length reached 90 minutes (orange).

**Figure 2 - Supplement 1.**
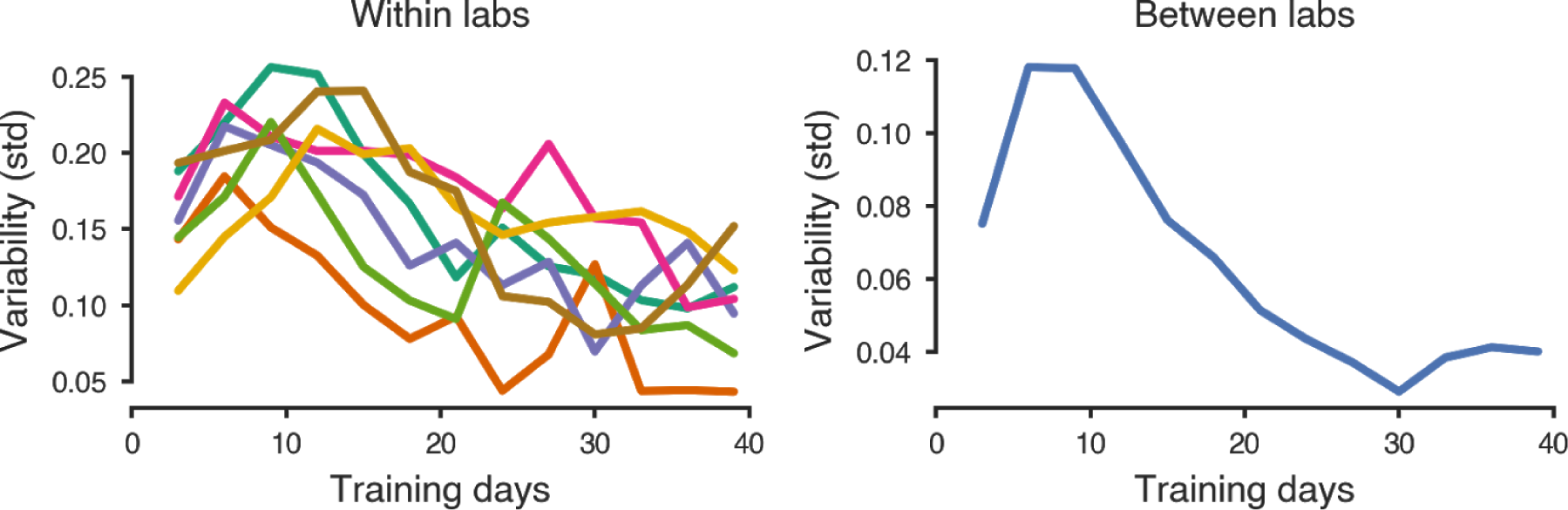
Performance variability within and across laboratories goes down over training time. **a-b**, Variability in performance (s.d. of % correct) in easy trials (100% and 50% contrast) (**a**) within (colors as in Figure 2, 3 and 4), and (**b**) across laboratories during the first 40 days of training.

**Figure 3 - Supplement 1.**
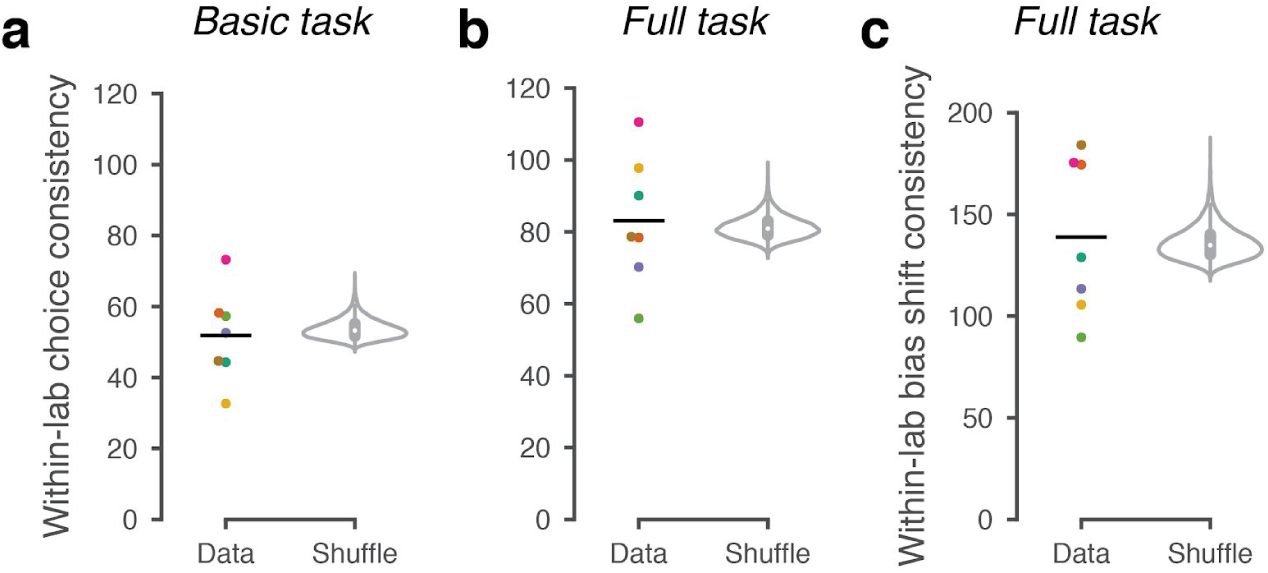
Within-lab consistency of mouse choices. We computed *within-lab choice consistency*, to measure the similarity in behavior across mice within an individual lab. (**a**) Within-lab choice consistency for the basic task (same data as in Figure 3). For each lab and each signed contrast we computed the variance across mice in the fraction of rightward choices. We then computed the inverse (consistency) and averaged the result across the signed contrasts (*dots*). We then averaged this measure across labs to quantify the overall within-lab consistency of choices (*horizontal line*). A null distribution was generated by randomly shuffling lab assignments between animals and computing the average within-lab choice variability 10,000 times (*violin plot*). If choice behavior were more consistent within labs than across labs, this shuffled measure would be lower than in the real data. Instead, for the basic task it was not lower than expected by chance (p = 0.73). (**b**) For the full task, we computed the choice fractions separately per signed contrast as well as prior block, before taking the average consistency (1/variance) across mice. Within-lab choice consistency on the full task (same data as in Figure 4) was not higher than expected by chance, p = 0.25. Notice that choice consistency is higher on the full task than the basic task; this likely reflects both increased training on the task, and a stronger constraint on choice behavior through the full task’s block structure. (**c**) As in a, b, but measuring the within-lab consistency of ‘bias shift’ between the 20:80 and 80:20 blocks (as in Figure 4d,e). Within-lab consistency in bias shift was not higher than expected by chance (p = 0.31).

**Figure 3 - Supplement 2.**
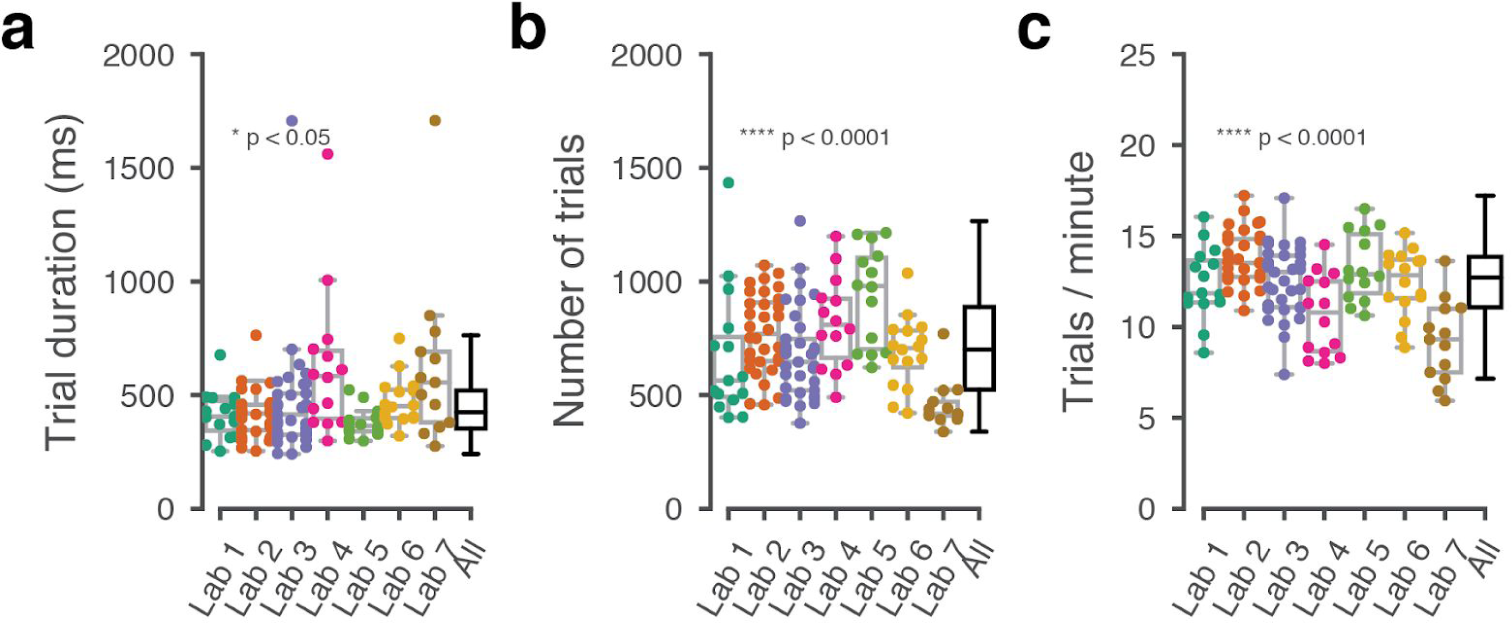
Behavioral metrics that were not explicitly trained, varied over labs. **a-b**, For the three sessions at which a mouse achieved basic task proficiency, box plots showing the distribution of (**a**) trial duration from go cue to correct or incorrect outcome, (**b**) the average number of trials in each of the three sessions, (**c**) number of trials per minute.

**Figure 3 - Supplement 3.**
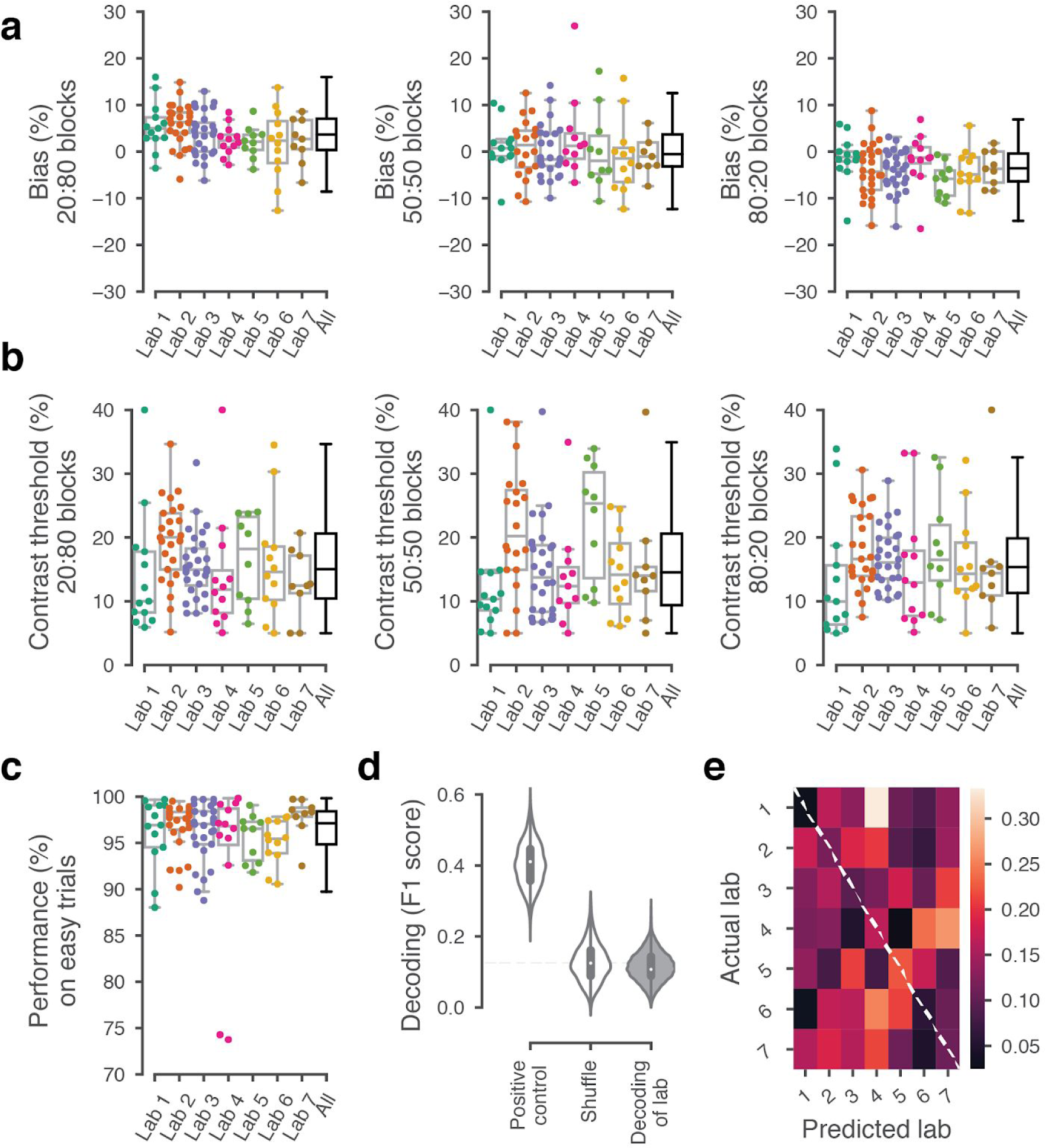
Behavior was indistinguishable over labs for the three sessions after first performing the full task. For the first 3 sessions of performing the full task (triggered by achieving proficiency in the basic task, defined by a specific set of criteria, Figure 1 - Supplement 1d), **a** Bias for each block prior did not vary significantly over labs (Kruskal-Wallis test, 20:80 blocks; p = 0.96, 50:50 block; p = 0.96, 80:20 block; p = 0.89) **b** The contrast thresholds also did not vary systematically over labs (Kruskal-Wallis test, 20:80 block; p = 0.078, 50:50 block; p = 0.12, 80:20 block; p = 0.17), **c** Performance on 100% contrast trials neither (Kruskal-Wallis test, p = 0.15), **d** The between-lab Naive Bayes classifier trained on the data in **a-c** did not perform above chance level when trying to predict the lab membership of mice. **e** Normalized confusion matrix for the classifier in **d**.

**Figure 3 - Supplement 4.**
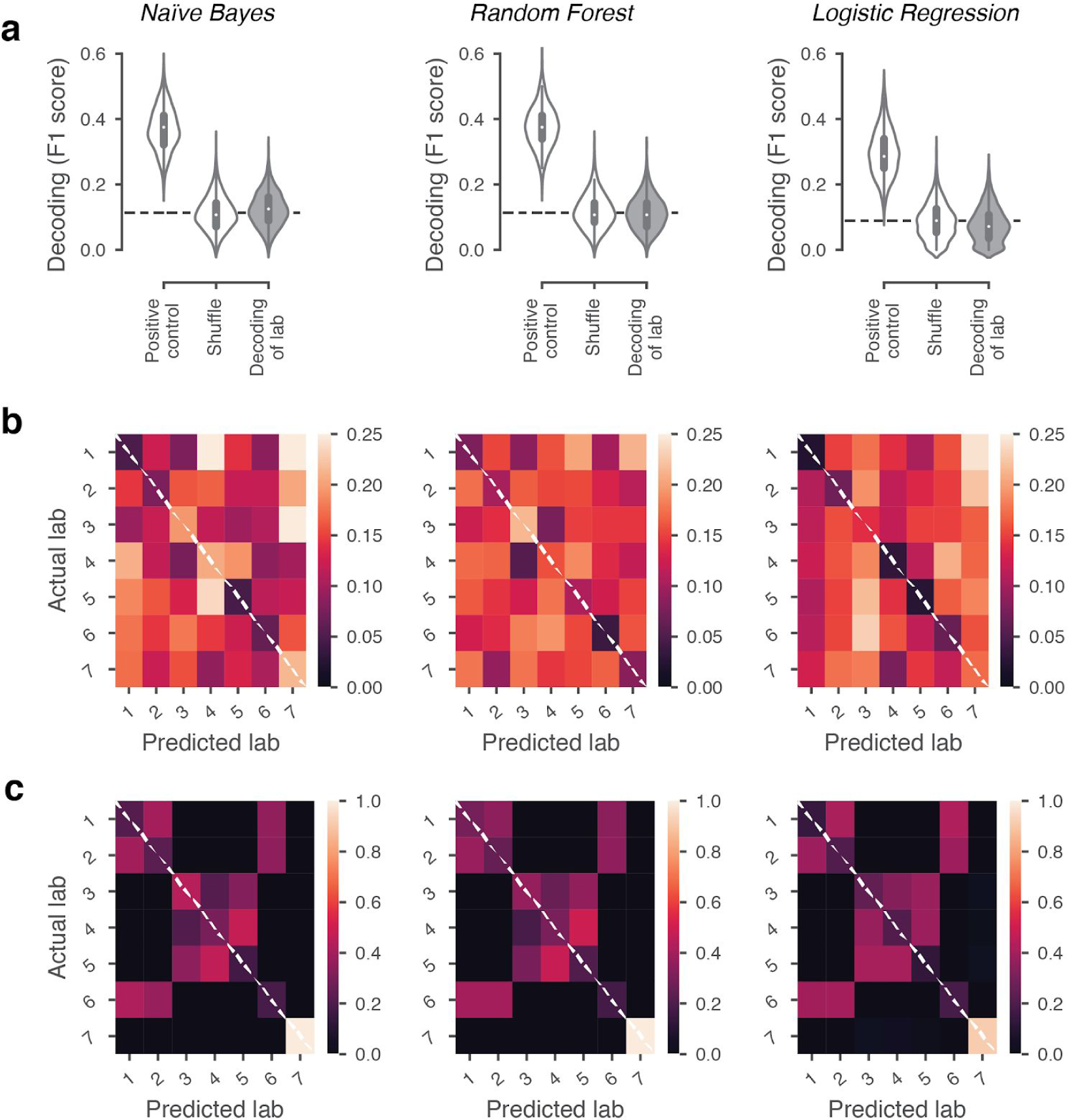
Classification of lab membership by three different classifiers could not predict lab membership from behavior. **a**, Classification performance of a Naive Bayes, Random Forest and Logistic Regression classifier while predicting lab membership based on behavioral metrics from Figure 3c-e like in Figure 3f. From each lab, 8 mice were randomly subsampled 2,000 times and for each iteration lab membership was classified using leave-one-out cross-validation. The positive control included the time zone in which a mouse was trained in the dataset and in the shuffle condition the lab labels were shuffled. **b**, Normalized confusion matrices for the classifiers in **a** which indicates the proportion of occurrences that a mouse was classified to be in the predicted lab (x-axis) while it was from the ‘actual lab’ (y-axis). **c**, Normalized confusion matrices as in **b**, for the positive control. Labs that are located in the same time zone from clear clusters, Lab 7 is always correctly predicted because it’s the only lab that doesn’t have any other labs in the same time zone.

**Figure 5 - Supplement 1.**
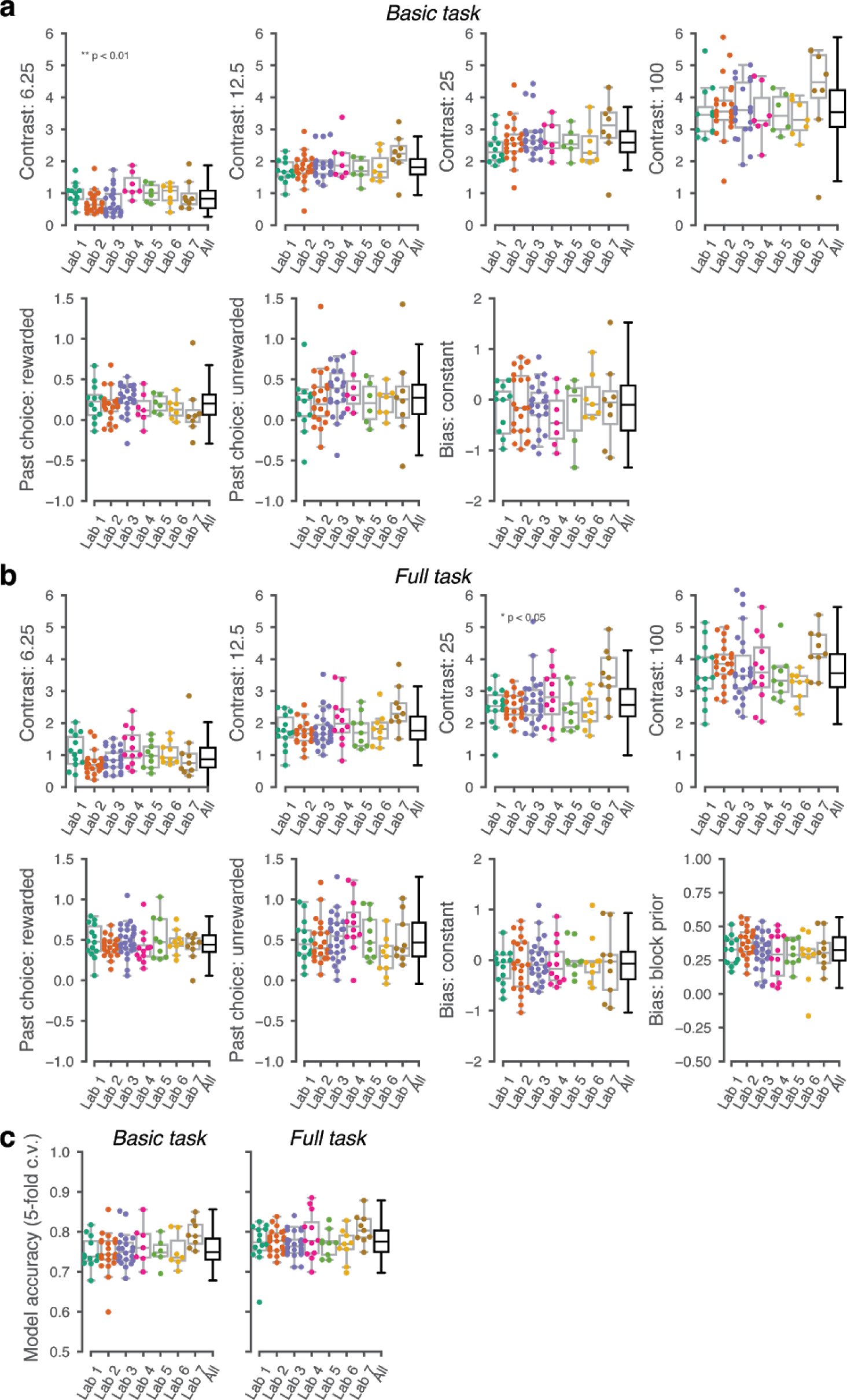
GLM accuracy across labs. Cross validated GLM accuracy (threshold: 50%) across mice and laborations in the **(a)** basic and **(b)** full task. Each point represents the average accuracy for each mouse.

**Figure 5 - Supplement 2.**
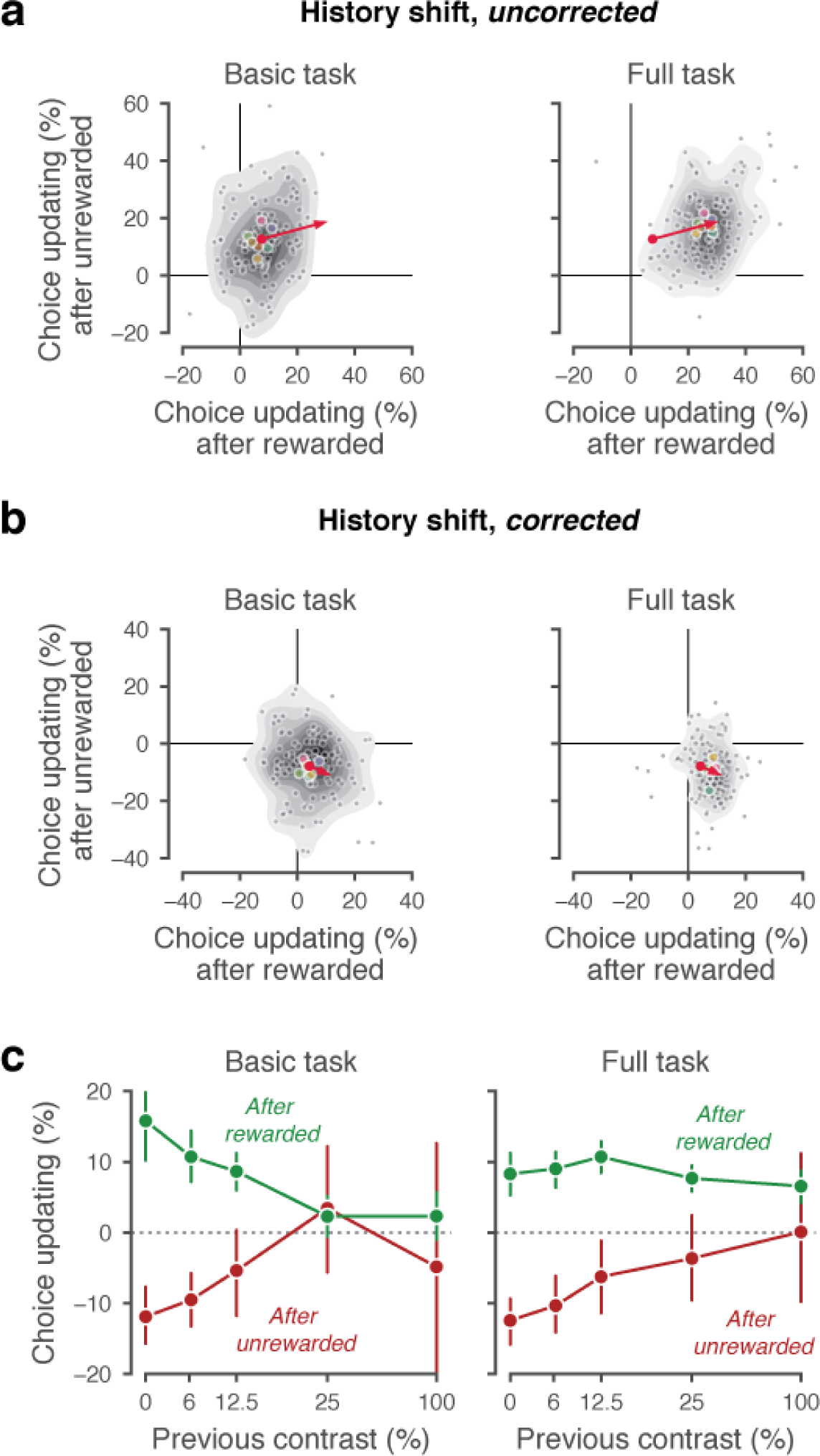
History-dependent choice updating. (**a**) Representing each animal’s ‘history strategy’, defined as the bias shift in their psychometric function as a function of the choice made on the previous trial, separately for when this trial was rewarded or unrewarded. Each animal is shown as a dot, with lab-averages shown larger colored dots. Contours indicate a two-dimensional kernel density estimate across all animals. The red arrow shows the group average in the basic task at its origin, and in the full task at its end (replicated between the left and right panel). (**b**) as a, but with the strategy space corrected for slow fluctuations in decision bound (Lak et al., 2020a). When taking these slow state-changes into account, the majority of animals use a win-stay lose-switch strategy. (**c**) History-dependent choice updating, after removing the effect of slow fluctuations in decision bound, as a function of the previous trial’s reward and visual contrast. After rewarded trials, choice updating is largest when the visual stimulus was highly uncertain (i.e. had low contrast) but strongly diminished after more certain, rewarded trials. This is in line with predictions from Bayesian models, where an agent continually updates its beliefs about the upcoming stimuli with sensory evidence (Lak et al., 2020a; Mendonça et al., 2018)).

## Notes

https://data.internationalbrainlab.org

https://github.com/int-brain-lab

## References

Aoki, R., Tsubota, T., Goya, Y., and Benucci, A. (2017). An automated platform for high-throughput mouse behavior and physiology with voluntary head-fixation. Nat. Commun.8a, 1–9.

Ashwood, Z., et al., IBL Collaboration, and Pillow, J.W. (2019). State-dependent modeling of psychophysical behavior during decision making. Soc. Neurosci. 2019 Online.

Bak, J.H., Choi, J.Y., Akrami, A., Witten, I., and Pillow, J.W. (2016). Adaptive optimal training of animal behavior. In Advances in Neural Information Processing Systems 29, D.D. Lee, M. Sugiyama, U.V. Luxburg, I. Guyon, and R. Garnett, eds. (Curran Associates, Inc.), pp. 1947–1955.

Baker, M. (2016). 1,500 scientists lift the lid on reproducibility. Nat. News 533, 452.

Beraldo, F.H., Palmer, D., Memar, S., Wasserman, D.I., Lee, W.-J.V., Liang, S., Creighton, S.D., Kolisnyk, B., Cowan, M.F., Mels, J., et al. (2019). MouseBytes, an open-access high-throughput pipeline and database for rodent touchscreen-based cognitive assessment. ELife 8, e49630.

Botvinik-Nezer, R., Holzmeister, F., Camerer, C.F., Dreber, A., Huber, J., Johannesson, M., Kirchler, M., Iwanir, R., Mumford, J.A., Adcock, R.A., et al. (2020). Variability in the analysis of a single neuroimaging dataset by many teams. Nature 582, 84–88.

Brand, A., Allen, L., Altman, M., Hlava, M., and Scott, J. (2015). Beyond authorship: attribution, contribution, collaboration, and credit. Learn. Publ.28, 151–155.

Burgess, C.P., Lak, A., Steinmetz, N.A., Zatka-Haas, P., Reddy, C.B., Jacobs, E.A.K., Linden, J.F., Paton, J.J., Ranson, A., Schröder, S., et al. (2017). High-Yield Methods for Accurate Two-Alternative Visual Psychophysics in Head-Fixed Mice. Cell Rep.20, 2513–2524.

Busse, L., Ayaz, A., Dhruv, N.T., Katzner, S., Saleem, A.B., Schölvinck, M.L., Zaharia, A.D., and Carandini, M. (2011). The detection of visual contrast in the behaving mouse. J. Neurosci. Off. J. Soc. Neurosci.31, 11351–11361.

Button, K.S., Ioannidis, J.P.A., Mokrysz, C., Nosek, B.A., Flint, J., Robinson, E.S.J., and Munafò, M.R. (2013). Power failure: why small sample size undermines the reliability of neuroscience. Nat. Rev. Neurosci.14, 365–376.

Bycroft, C., Freeman, C., Petkova, D., Band, G., Elliott, L.T., Sharp, K., Motyer, A., Vukcevic, D., Delaneau, O., O’Connell, J., et al. (2018). The UK Biobank resource with deep phenotyping and genomic data. Nature 562, 203–209.

Camerer, C.F., Dreber, A., Holzmeister, F., Ho, T.-H., Huber, J., Johannesson, M., Kirchler, M., Nave, G., Nosek, B.A., Pfeiffer, T., et al. (2018). Evaluating the replicability of social science experiments in Nature and Science between 2010 and 2015. Nat. Hum. Behav.2, 637–644.

Carandini, M., and Churchland, A.K. (2013). Probing perceptual decisions in rodents. Nat Neurosci 16, 824–831.

CERN Education, Communications and Outreach Group (2018). CERN Annual Report 2017 (CERN).

Charles, A.S., Falk, B., Turner, N., Pereira, T.D., Tward, D., Pedigo, B.D., Chung, J., Burns, R., Ghosh, S.S., Kebschull, J.M., et al. (2020). Toward Community-Driven Big Open Brain Science: Open Big Data and Tools for Structure, Function, and Genetics. Annu. Rev. Neurosci.43, null.

Chesler, E.J., Wilson, S.G., Lariviere, W.R., Rodriguez-Zas, S.L., and Mogil, J.S. (2002). Influences of laboratory environment on behavior. Nat. Neurosci.5, 1101–1102.

Cohen, M.R., and Maunsell, J.H.R. (2009). Attention improves performance primarily by reducing interneuronal correlations. Nat. Neurosci.12, 1594–1600.

Corrado, G.S., Sugrue, L.P., Seung, H.S., and Newsome, W.T. (2005). Linear-Nonlinear-Poisson models of primate choice dynamics. J. Exp. Anal. Behav.84, 581–617.

Crabbe, J.C., Wahlsten, D., and Dudek, B.C. (1999). Genetics of Mouse Behavior: Interactions with Laboratory Environment. Science 284, 1670–1672.

Dickinson, M.E., Flenniken, A.M., Ji, X., Teboul, L., Wong, M.D., White, J.K., Meehan, T.F., Weninger, W.J., Westerberg, H., Adissu, H., et al. (2016). High-throughput discovery of novel developmental phenotypes. Nature 537, 508–514.

Fan, Y., Gold, J.I., and Ding, L. (2018). Ongoing, rational calibration of reward-driven perceptual biases. ELife 7, e36018.

Fish, V.L., Akiyama, K., Bouman, K.L., Chael, A.A., Johnson, M.D., Doeleman, S.S., Blackburn, L., Wardle, J.F.C., Freeman, W.T., and The Event Horizon Telescope Collaboration (2016). Observing—and Imaging—Active Galactic Nuclei with the Event Horizon Telescope. Galaxies 4, 54.

Forscher, P.S., Wagenmakers, E.-J., Coles, N.A., Silan, M.A., and IJzerman, H. (2020). A Manifesto for Team Science (PsyArXiv).

Frank, M.C., Bergelson, E., Bergmann, C., Cristia, A., Floccia, C., Gervain, J., Hamlin, J.K., Hannon, E.E., Kline, M., Levelt, C., et al. (2017). A Collaborative Approach to Infant Research: Promoting Reproducibility, Best Practices, and Theory-Building. Infancy Off. J. Int. Soc. Infant Stud.22, 421–435.

Glickfeld, L.L., Reid, R.C., and Andermann, M.L. (2014). A mouse model of higher visual cortical function. Curr. Opin. Neurobiol.24, 28–33.

Guo, Z.V., Hires, S.A., Li, N., O’Connor, D.H., Komiyama, T., Ophir, E., Huber, D., Bonardi, C., Morandell, K., Gutnisky, D., et al. (2014). Procedures for Behavioral Experiments in Head-Fixed Mice. PLOS ONE 9, e88678.

Harris, C.R., Millman, K.J., van der Walt, S.J., Gommers, R., Virtanen, P., Cournapeau, D., Wieser, E., Taylor, J., Berg, S., Smith, N.J., et al. (2020). Array programming with NumPy. Nature 585, 357–362.

Herrnstein, R.J. (1961). Relative and absolute strength of response as a function of frequency of reinforcement,. J. Exp. Anal. Behav.4, 267–272.

International Brain Laboratory, Bonacchi, N., Chapuis, G., Churchland, A., Harris, K.D., Rossant, C., Sasaki, M., Shen, S., Steinmetz, N.A., Walker, E.Y., et al. (2019). Data architecture and visualization for a large-scale neuroscience collaboration. BioRxiv 827873.

Ioannidis, J.P.A. (2005). Why Most Published Research Findings Are False. PLoS Med. 2.

Kafkafi, N., Agassi, J., Chesler, E.J., Crabbe, J.C., Crusio, W.E., Eilam, D., Gerlai, R., Golani, I., Gomez-Marin, A., Heller, R., et al. (2018). Reproducibility and replicability of rodent phenotyping in preclinical studies. Neurosci. Biobehav. Rev.87, 218–232.

Koscielny, G., Yaikhom, G., Iyer, V., Meehan, T.F., Morgan, H., Atienza-Herrero, J., Blake, A., Chen, C.-K., Easty, R., Di Fenza, A., et al. (2014). The International Mouse Phenotyping Consortium Web Portal, a unified point of access for knockout mice and related phenotyping data. Nucleic Acids Res.42, D802–809.

Lak, A., Hueske, E., Hirokawa, J., Masset, P., Ott, T., Urai, A.E., Donner, T.H., Carandini, M., Tonegawa, S., Uchida, N., et al. (2020a). Reinforcement biases subsequent perceptual decisions when confidence is low, a widespread behavioral phenomenon. ELife 9, e49834.

Lak, A., Okun, M., Moss, M.M., Gurnani, H., Farrell, K., Wells, M.J., Reddy, C.B., Kepecs, A., Harris, K.D., and Carandini, M. (2020b). Dopaminergic and Prefrontal Basis of Learning from Sensory Confidence and Reward Value. Neuron 105, 700-711.e6.

Lau, B., and Glimcher, P.W. (2005). Dynamic Response-by-Response Models of Matching Behavior in Rhesus Monkeys. J. Exp. Anal. Behav.84, 555–579.

Lopes, G., Bonacchi, N., Frazão, J., Neto, J.P., Atallah, B.V., Soares, S., Moreira, L., Matias, S., Itskov, P.M., Correia, P.A., et al. (2015). Bonsai: an event-based framework for processing and controlling data streams. Front. Neuroinformatics 9.

Lopes, G., Farrell, K., Horrocks, E.A.B., Lee, C.-Y., Morimoto, M.M., Muzzu, T., Papanikolaou, A., Rodrigues, F.R., Wheatcroft, T., Zucca, S., et al. (2020). BonVision – an open-source software to create and control visual environments. BioRxiv 2020.03.09.983775.

Makel, M.C., Plucker, J.A., and Hegarty, B. (2012). Replications in Psychology Research: How Often Do They Really Occur? Perspect. Psychol. Sci. J. Assoc. Psychol. Sci.7, 537–542.

Mathis, A., Mamidanna, P., Cury, K.M., Abe, T., Murthy, V.N., Mathis, M.W., and Bethge, M. (2018). DeepLabCut: markerless pose estimation of user-defined body parts with deep learning. Nat Neurosci 21, 1281–1289.

McGinley, M.J., Vinck, M., Reimer, J., Batista-Brito, R., Zagha, E., Cadwell, C.R., Tolias, A.S., Cardin, J.A., and McCormick, D.A. (2015). Waking State: Rapid Variations Modulate Neural and Behavioral Responses. Neuron 87, 1143–1161.

Mendonça, A.G., Drugowitsch, J., Vicente, M.I., DeWitt, E., Pouget, A., and Mainen, Z.F. (2018). The impact of learning on perceptual decisions and its implication for speed-accuracy tradeoffs. BioRxiv 501858.

Miller, K.J., Botvinick, M.M., and Brody, C.D. (2019). From predictive models to cognitive models: An analysis of rat behavior in the two-armed bandit task. BioRxiv 461129.

Norton, E.H., Acerbi, L., Ma, W.J., and Landy, M.S. (2019). Human online adaptation to changes in prior probability. PLOS Comput. Biol.15, e1006681.

O’Connor, D.H., Huber, D., and Svoboda, K. (2009). Reverse engineering the mouse brain. Nature 461, 923–929.

Pedregosa, F., Varoquaux, G., Gramfort, A., Michel, V., Thirion, B., Grisel, O., Blondel, M., Prettenhofer, P., Weiss, R., Dubourg, V., et al. (2011). Scikit-learn: Machine Learning in Python. J. Mach. Learn. Res.12, 2825–2830.

Pinto, L., Koay, S.A., Engelhard, B., Yoon, A.M., Deverett, B., Thiberge, S.Y., Witten, I.B., Tank, D.W., and Brody, C.D. (2018). An Accumulation-of-Evidence Task Using Visual Pulses for Mice Navigating in Virtual Reality. Front. Behav. Neurosci.12.

Pisupati, S., Chartarifsky-Lynn, L., Khanal, A., and Churchland, A.K. (2019). Lapses in perceptual decisions reflect exploration (Neuroscience).

Poddar, R., Kawai, R., and Ölveczky, B.P. (2013). A Fully Automated High-Throughput Training System for Rodents. PLOS ONE 8, e83171.

Poldrack, R.A., and Gorgolewski, K.J. (2014). Making big data open: data sharing in neuroimaging. Nat. Neurosci.17, 1510–1517.

Reback, J., McKinney, W., jbrockmendel, Bossche, J.V. den, Augspurger, T., Cloud, P., gfyoung, Sinhrks, Klein, A., Roeschke, M., et al. (2020). pandas-dev/pandas: Pandas 1.0.1 (Zenodo).

Roy, N.A., Bak, J.H., Akrami, A., Brody, C.D., and Pillow, J.W. (2020). Extracting the Dynamics of Behavior in Decision-Making Experiments. BioRxiv 2020.05.21.109678.

Scott, B.B., Constantinople, C.M., Erlich, J.C., Tank, D.W., and Brody, C.D. (2015). Sources of noise during accumulation of evidence in unrestrained and voluntarily head-restrained rats. ELife 4, e11308.

Seabold, S., and Perktold, J. (2010). Statsmodels: Econometric and Statistical Modeling with Python. p. Smith, N.J., Hudon, C., broessli, Skipper Seabold, Peter Quackenbush, Michael Hudson-Doyle, Max

Humber, Katrin Leinweber, Hassan Kibirige, Cameron Davidson-Pilon, et al. (2018). pydata/patsy: v0.5.1 (Zenodo).

Sorge, R.E., Martin, L.J., Isbester, K.A., Sotocinal, S.G., Rosen, S., Tuttle, A.H., Wieskopf, J.S., Acland, E.L., Dokova, A., Kadoura, B., et al. (2014). Olfactory exposure to males, including men, causes stress and related analgesia in rodents. Nat. Methods 11, 629–632.

Tanner Jr., W.P., and Swets, J.A. (1954). A decision-making theory of visual detection. Psychol. Rev.61, 401–409.

The H.E.S.S. Galactic plane survey (2018). The H.E.S.S. Galactic plane survey. Astron. Astrophys. 612, A1.

The International Brain Laboratory (2017). An International Laboratory for Systems and Computational Neuroscience. Neuron 96, 1213–1218.

Tuttle, A.H., Philip, V.M., Chesler, E.J., and Mogil, J.S. (2018). Comparing phenotypic variation between inbred and outbred mice. Nat. Methods 15, 994–996.

Urai, A.E., Braun, A., and Donner, T.H. (2017). Pupil-linked arousal is driven by decision uncertainty and alters serial choice bias. Nat. Commun.8, 14637.

Urai, A.E., Aguillon-Rodriguez, V., Laranjeira, I.C., Cazettes, F., Laboratory, T.I.B., Mainen, Z.F., and Churchland, A.K. (2020). Citric Acid Water as an Alternative to Water Restriction for High-Yield Mouse Behavior. BioRxiv 2020.03.02.973016.

de Vries, S.E.J., Lecoq, J.A., Buice, M.A., Groblewski, P.A., Ocker, G.K., Oliver, M., Feng, D., Cain, N., Ledochowitsch, P., Millman, D., et al. (2020). A large-scale standardized physiological survey reveals functional organization of the mouse visual cortex. Nat. Neurosci.23, 138–151.

Waskom, M., Botvinnik, O., O’Kane, D., Hobson, P., Lukauskas, S., Gemperline, D.C., Augspurger, T., Halchenko, Y., Cole, J.B., Warmenhoven, J., et al. (2017). mwaskom/seaborn: v0.8.1 (September 2017) (Zenodo).

Whiteley, L., and Sahani, M. (2008). Implicit knowledge of visual uncertainty guides decisions with asymmetric outcomes. J. Vis.8, 2–2.

Wool, L., and The International Brain Laboratory. (2020). PsyArXiv Preprints | Knowledge across networks: how to build a global neuroscience collaboration. PsyArxiv.

