## Appendix 1 for "Standardized and reproducible measurement of decision-making in mice"

INTERNATIONAL BRAIN LABORATORY

### Appendix 1: IBL protocol for headbar implant surgery in mice

**The latest version of this protocol can be found on Figshare via this specific link.**

All IBL protocols cited in the article can be found on Figshare via this master link.

The latest version of the CAD models and technical drawings for the manufactured rig parts can be found on Figshare via this specific link.

The latest version of the whole rig CAD assembly can be found on Figshare via this specific link.

**Please cite the associated article** when using any of these materials and protocols:

The International Brain Laboratory et al. (2020) *Standardized and reproducible measurement of decision-making in mice.* bioRxiv, 909838. https://doi.org/10.1101/2020.01.17.909838
